# Structural and functional properties of a plant NRAMP related aluminum transporter

**DOI:** 10.1101/2022.12.21.521437

**Authors:** Karthik Ramanadane, Márton Liziczai, Dragana Markovic, Monique S. Straub, Gian T. Rosalen, Anto Udovcic, Raimund Dutzler, Cristina Manatschal

## Abstract

The transport of transition metal ions by members of the SLC11/NRAMP family constitutes a ubiquitous mechanism for the uptake of Fe^2+^ and Mn^2+^ across all kingdoms of life. Despite the strong conservation of the family, two of its branches have evolved a distinct substrate preference with one mediating Mg^2+^ uptake in prokaryotes and another the transport of Al^3+^ into plant cells. Our previous work on the SLC11 transporter from *Eggerthella lenta* revealed the basis for its Mg^2+^ selectivity (Ramanadane et al., 2022). Here we have addressed the structural and functional properties of a putative Al^3+^ transporter from *Setaria italica.* We show that the protein transports diverse divalent metal ions and binds the trivalent ions Al^3+^ and Ga^3+^, which are both presumable substrates. Its cryo-EM structure displays an occluded conformation that is closer to an inward-than an outward-facing state with a binding site that is remodeled to accommodate the increased charge density of its transported substrate.

## Introduction

Owing to their unique properties to form coordinative interactions with elements containing free electron pairs and their ability to readily change their oxidation states, Fe^2+^ and Mn^2+^ are important trace elements in biology. The uptake of both ions into cells requires an elaborate mechanism that allows their selection from a large background of alkaline earth metal ions and their efficient concentration within cells. This formidable task is performed by members of the SLC11 family, which constitute a conserved family of secondary active transporters that facilitate the cotransport of transition metal ions together with protons acting as energy source (Courville et al., 2006; Gunshin et al., 1997; Montalbetti et al., 2013). To be able to do this across all kingdoms of life, the common molecular features conferring ion selectivity and proton coupling are strongly conserved within different branches of the family (Bozzi and Gaudet, 2021). However, despite this strong conservation, there are two distinct clades of the family, which have evolved a different substrate selectivity. These comprise the large group of NRAMP related Mg^2+^ transporters (NRMTs) and a smaller group of NRAMP related Al^3+^ transporters (NRATs) (Chauhan et al., 2021; Shin et al., 2014; Xia et al., 2010). In a recent study, we have shown how NRMTs have adapted their molecular architecture to facilitate Mg^2+^ uptake, by characterizing a transporter from the bacterium *Eggerthella lenta* termed EleNRMT (Ramanadane et al., 2022). Due to its small ionic radius, the stripping of the interacting solvent is particularly costly for Mg^2+^, and its transport by EleNRMT thus proceeds as hydrated ion retaining most of its first coordination shell. Besides Mg^2+^, the protein also transports Mn^2+^ but not Ca^2+^. In NRMTs, several of the residues coordinating ions in transition metal transporters of the family are exchanged, including a conserved methionine, which forms coordinative interactions with transition metals while preventing the binding of alkaline earth metal ions (Bozzi et al., 2016). The reduction of the side chain volume of interacting residues in combination with small rearrangements of the backbone have increased the volume of the binding site to accommodate the hydrated ion (Ramanadane et al., 2022). Unlike in proton-coupled family members, transport in EleNRMT is passive with residues that have been assigned to participate in H^+^ transport being replaced by hydrophobic side chains (Ramanadane et al., 2022).

Besides the prokaryotic NRMTs, which constitute a large branch of the family that is distantly related to transition metal ion transporters of the same family, the NRATs form a small clade in plants, whose evolutionary relationship to classical SLC11 transporters is much closer (Ramanadane et al., 2022). These proteins have evolved to combat Al^3+^ toxicity in soil (Chauhan et al., 2021). As strong Lewis acid, this trivalent cation strongly interacts with membranes to compromise their integrity. As one of the mechanisms to neutralize the adverse effect of Al^3+^, NRATs were proposed to catalyze its uptake into the cytoplasm of root cells acting in concert with other proteins, which mediate the further transport into vacuoles where it is neutralized by the complexation with dicarboxylates (Xia et al., 2010).

To clarify the mechanism by which NRATs transport trivalent cations, we have here studied the function and structure of a transporter from *Setaria italica* (termed SiNRAT). We show that this protein broadly transports divalent transition metal ions by a mechanism that is not coupled to protons. Although, for technical reasons, transport of trivalent cations could not directly be assayed, we provide evidence for their specific interactions with the binding site thus defining them as likely substrates. The cryo-EM structure of SiNRAT defines a protein that shares the hallmarks of the SLC11 family. It resides in an occluded conformation where the access to the binding site is sealed from both sides of the membrane. This structure harbors an ion binding region for Al^3+^ located inside an aqueous pocket, which contains the predicted replacements of coordinating residues and additionally an acidic amino acid that increases the negative charge density of the site to accommodate the larger net charge of the transported substrate.

## Results

### Functional characterization of SiNRAT

In the course of a previous phylogenetic analysis, we were able to spot five homologues of the prototypic NRAT1 from *Oryza sativa* (XP_015625418.1, Uniprot: Q6ZG85) in Blast searches of eukaryotic sequence databases (Ramanadane et al., 2022). All identified proteins contain the described hallmarks of NRATs, which distinguishes this clade of the SLC11 family from the bulk of transition metal ion transporters. Differences include variations in the putative ion coordinating site, where the conserved DPGN motif on α1 of classical NRAMPs is changed to DPSN and the A/CxxM motif on α6 to AxxT (Lu et al., 2018; Ramanadane et al., 2022). Latter replaces the conserved methionine, whose sulfur atom is involved in transition metal ion interactions, with a threonine, which offers a hard oxygen ligand for metal ion interactions. To identify proteins that are suitable for structural and functional investigations, we have generated constructs of all five NRAT homologs, each containing a C-terminal fusion of eGFP and an SBP tag used for affinity purification preceded by a 3C protease cleavage site. We have subsequently investigated their expression upon transient transfection of HEK293 cells followed by an initial biochemical characterization by fluorescence size exclusion chromatography (Hattori et al., 2012). These experiments have singled out the protein from *Setaria italica* (termed SiNRAT, XP_004952002.1) as construct with comparably high expression level and promising biochemical behavior. We have subsequently scaled up the expression of SiNRAT in suspension culture and purified it in the detergent n-dodecyl-β-D-maltopyranoside (DDM) for further studies (Figure 1—figure supplement 1A, B).

For a characterization of its transport properties, SiNRAT was reconstituted into proteoliposomes and subjected to previously established fluorescence-based assays that have permitted the functional investigation of other members of the SLC11 family (Ehrnstorfer et al., 2014; Ehrnstorfer et al., 2017; Ramanadane et al., 2022). We have initially studied the transport of Mn^2+^, which is a common substrate of SLC11 transporters that can be monitored using the fluorophore calcein. This assay revealed a concentration-dependent quenching of calcein as a consequence of the influx of Mn^2+^ into the proteoliposomes (Figure 1A). Transport is strongly dependent on the Mn^2+^ concentration and saturates with an apparent K_M_ of 13 μM reflecting the binding of the transported substrate to a saturable site (Figure 1A, B). We subsequently investigated impairment of Mn^2+^ transport by the alkaline earth metal ions Ca^2+^ and Mg^2+^ and found a concentration-dependent competition by both ions, which is indicative for their specific binding to the protein (Figure 1C, D, Figure 1—figure supplement 1C, D). To investigate whether both ions would also be transported, we directly monitored their influx into proteoliposomes employing the fluorophores Fura-2 for the detection of Ca^2+^ and Magnesium green for Mg^2+^. In both cases we found a weak but concentration-dependent signal that reflects uptake of both ions (Figure 1E, F, Figure 1—figure supplement 1E, F).

**Figure 1.**
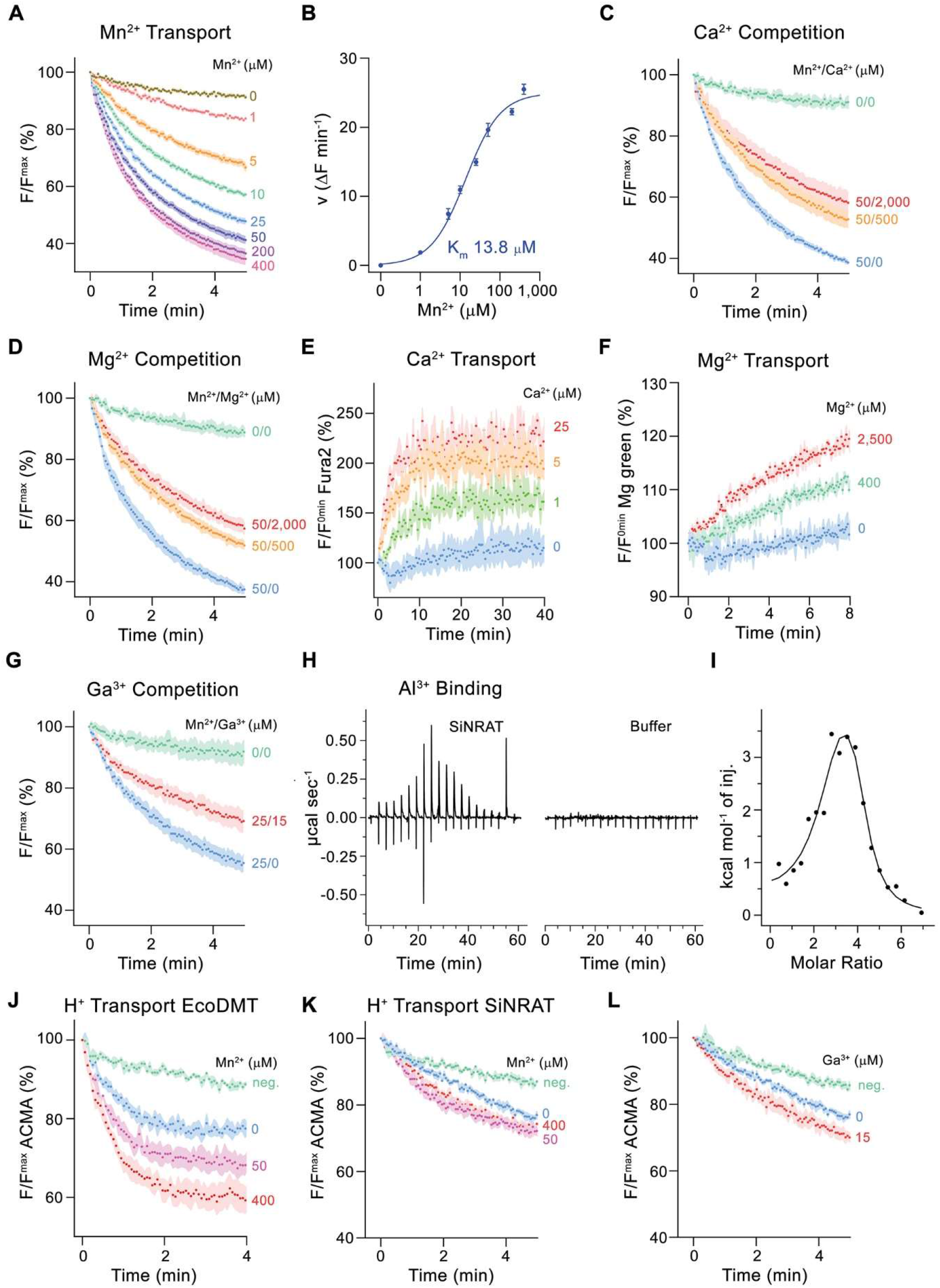
Functional characterization of SiNRAT. (**A**) SiNRAT mediated Mn^2+^ transport into proteoliposomes (6 experiments from 4 independent reconstitutions). (**B**) Mn^2+^ concentration dependence of transport. Initial velocities were derived from individual traces of experiments displayed in **A**, the solid line shows the fit to a Michaelis–Menten equation with an apparent *K*_m_ of 13.8 μM. (**C**) Mn^2+^ transport in presence of Ca^2+^ (3 experiments from 3 independent reconstitutions for all conditions). (**D**) Mn^2+^ transport in presence of Mg^2+^ (3 experiments from 3 independent reconstitutions for all conditions). (**E**) SiNRAT mediated Ca^2+^ import into proteoliposomes assayed with the fluorophore Fura2 trapped inside the liposome (3 experiments from 3 independent reconstitutions for all conditions). (**F**) Assay of Mg^2+^ import into proteoliposomes assayed with the fluorophore Magnesium green (3 experiments from 3 independent reconstitutions for all conditions). (**G**) Mn^2+^ transport in presence of Ga^3+^ (5 experiments from 2 independent reconstitutions for conditions containing Mn^2+^, 4 experiments for the condition 0 μM Mn^2+^ / 0 μM Ga^3+^). **A, C, D, G** Uptake of Mn^2+^ was assayed by the quenching of the fluorophore calcein trapped inside the vesicles. (**H**) Thermograms of Al^3+^ binding to SiNRAT (left) and buffer (right) obtained from isothermal titration calorimetry experiments. (**I)** Binding isotherm of Al^3+^ was fitted to a sum of two binding constants with the binding isotherm depicted as solid line. **J-L** Assay of H^+^ transport with the fluorophore ACMA. Experiments probing metal ion coupled H^+^ transport into proteoliposomes upon addition of metal ions to the outside. (**J**) H^+^ transport into proteoliposomes containing EcoDMT upon addition of Mn^2+^ (3 experiments from 2 independent reconstitutions for all conditions). H^+^ transport into proteoliposomes containing SiNRAT upon addition of (**K**) Mn^2+^ (4 experiments from 3 independent reconstitutions) and (**L**) Ga^3+^ (4 experiments from 3 independent reconstitutions for SiNRAT with no substrate and empty liposomes with 15 μM Ga^3+^ and 3 experiments from 3 independent reconstitutions for SiNRAT with 7.5 or 15 μM Ga^3+^). **A-G** and **J-L** Panels show mean of indicated number of experiments, errors are s.e.m.. **A**, **C**, **D-G**, **J-L**, Fluorescence is normalized to the value after addition of substrate (t=0). Applied ion concentrations are indicated. Negative controls (neg.) refer to empty liposomes in presence of 400 μM Mn^2+^ or 15 μM Ga^3+^, respectively. **Figure supplement 1.** Purification and assay data.

After demonstrating that SiNRAT is capable of mediating the transport of divalent metal ions with poor selectivity, we were interested in the transport properties of Al^3+^, which is the proposed physiological substrate. However, assaying this ion under *in vitro* conditions is complicated by the lack of suitable fluorophores and the fact that the chemical properties of Al^3+^, as strong Lewis acid with an ionic radius of 0.53 Å, leads to the destabilization of liposomes even at low μM concentrations. We thus turned our initial attention to the trivalent cation Ga^3+^, which is one period below Al^3+^ in the same group of the periodic system, whose larger ionic radius (0.62 Å) and consequent weaker Lewis acidity is less disruptive for lipid bilayers. In an assay to investigate the interference of Ga^3+^ with Mn^2+^ transport, we found an inhibition by Ga^3+^, thus emphasizing its interaction with the transporter (Figure 1G). At the same conditions, we did not observe such competition with the prokaryotic transition metal ion transporter EcoDMT, precluding the action of Ga^3+^ as non-specific inhibitor of the family (Figure 1—figure supplement 1G). Thus, despite the low solubility of Ga^3+^ under the assay conditions (Yu and Liao, 2011), which obscures a definitive assignment of its concentration dependence, our results suggest a specific effect of the ion on SiNRAT-mediated transport. Since equivalent studies with Al^3+^ are prevented by its disruptive properties on membrane integrity, we used isothermal titration calorimetry (ITC) to investigate the binding of the ion to SiNRAT in detergent solution. These experiments showed a specific signal upon Al^3+^ titration as a consequence of its interaction with the transporter (Figure 1H). However, at the neutral pH of the sample, the trivalent ion is at equilibrium with its hydroxyl complexes, which complicates data analysis (Kiss, 2013). The thermograms were thus best fitted to a sum of two binding constants, both in the low micromolar range (Figure 1I). Although precluding a definitive mechanistic interpretation, we use the ITC results of Al^3+^ binding as a qualitative assay of its interaction with SiNRAT and show later that the event is directly associated with its binding to the site that is relevant for ion transport.

We then investigated whether metal ion transport would be accompanied by a concentration-dependent acidification of vesicles, which is expected in case of coupled proton symport as observed for the prokaryotic SLC11 transporter EcoDMT (Figure 1J) (Ehrnstorfer et al., 2017). To this end, we have assayed the import of H^+^ at increasing concentrations of the transported ions Mn^2+^, Ca^2+^, Mg^2+^ and the putative substrate Ga^3+^ but did in no case detect evidence for metal ion concentrationdependent decrease of ACMA fluorescence, despite the negative membrane potential established by an outwardly directed K^+^ gradient, which would facilitate H^+^ transport (Figure 1K, L, Figure 1—figure supplement 1H, I). These findings suggest that, like in the Mg^2+^ transporter EleNRMT, SiNRAT acts as uncoupled transporter that facilitates the bidirectional transport of metal ions with poor selectivity.

### Structural characterization of SiNRAT

The described transport properties of SiNRAT are unique among characterized members of the SLC11 family. To define the underlying molecular features, we set out to determine its three-dimensional structure. Since the protein did not crystallize, we generated nanobody-based binders that, in complex with SiNRAT, increase the size of the protein sufficiently to permit its structure determination by cryo-electron microscopy (cryo-EM). These binders were obtained by immunization of alpacas with the purified protein and subsequently selected by phage display from a library generated from a blood sample (Pardon et al., 2014). The procedure allowed the identification of the nanobody Nb1^SiNRAT^ (short Nb1) as promising interaction partner that does not dissociate during size exclusion chromatography and which partly inhibits Mn^2+^ transport when added to the outside of liposomes (Figure 2—figure supplement 1, Table 2). For the SiNRAT-Nb1 complex, we have prepared samples for cryo-EM. Since data recorded in the detergent DDM did not permit a 3D reconstruction at high resolution, we have reconstituted the complex into amphipols (Kleinschmidt and Popot, 2014) and collected a large dataset, which ultimately yielded a cryo-EM density map at 3.66 Å (Figure 2—figure supplements 1-3, Table 1). This map was of high quality and allowed the unambiguous interpretation with an atomic model revealing details of the threedimensional organization of SiNRAT (Figure 2A, Figure 2—figure supplement 3).

**Figure 2.**
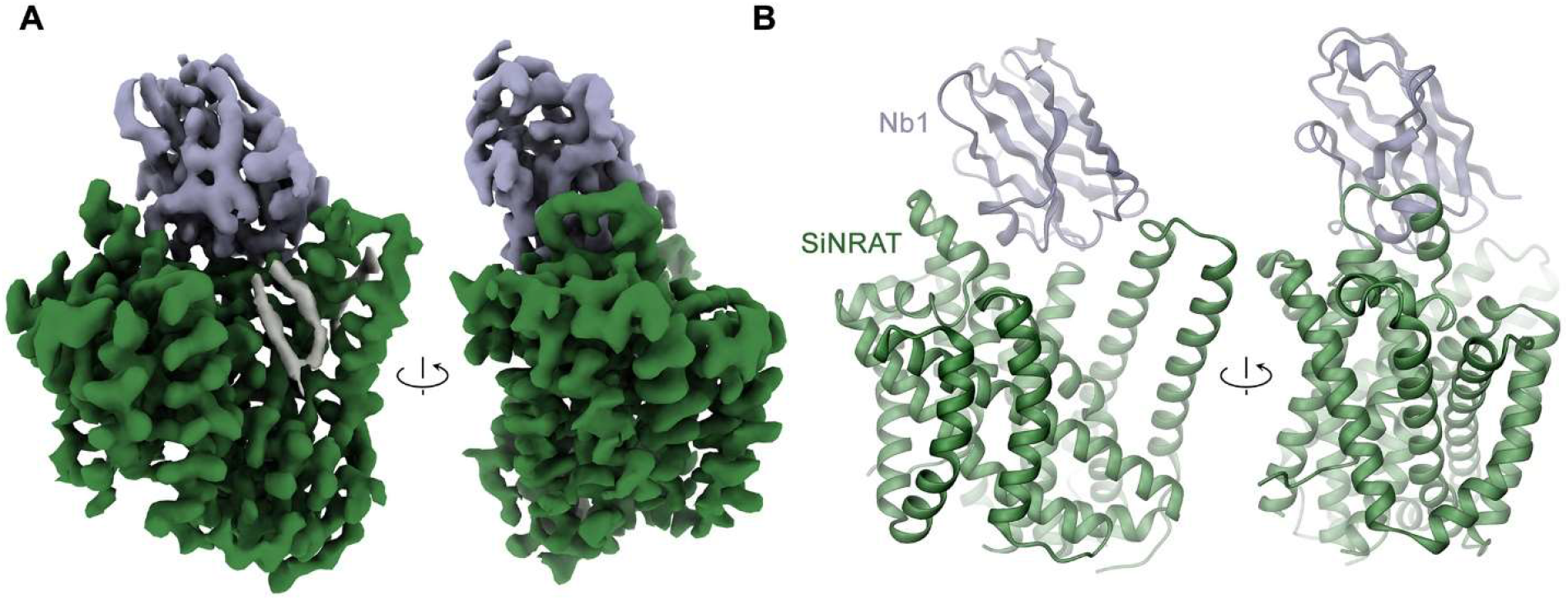
Structural characterization of the SiNRAT-Nb1 complex by Cryo-EM. (**A**) Cryo-EM density of SiNRAT in complex with Nb1 at 3.66 Å viewed from within the membrane at indicated orientations with the extracellular side on top and (**B**), ribbon representation of the complex in the same views. **A**, **B** Proteins are shown in unique colors, the density of a bound lipid in **A** is shown in grey. **Figure supplement 1.** Nanobody characterization. **Figure supplement 2.** Cryo-EM reconstitution of the SiNRAT-Nb1 complex. **Figure supplement 3.** Cryo-EM density of the SiNRAT-Nb1 complex. **Figure supplement 4.** SiNRAT-Nb1 interactions.

**Table 1.** Cryo-EM data collection, refinement and validation statistics.

|  |  |
| --- | --- |
|  | Dataset 1<br>SiNRAT-NB1 |
| **Data collection and processing** |  |
| Microscope | FEI Titan Krios |
| Camera | Gatan K3 GIF |
| Magnification | 130'000 |
| Voltage (kV) | 300 |
| Electron exposure (e<sup>-</sup>/Å<sup>2</sup>) | 69.739/ 69.63 |
| Defocus range (µm) | -1.0 to -2.4 |
| Pixel size* (Å) | 0.651 (0.3255) |
| Initial number of micrographs (no.) | 16530 |
| Initial particle images (no.) | 5,445,873 |
| Final particle images (no.) | 294,761 |
| Symmetry imposed | C1 |
| Map resolution (Å)<br>FSC threshold | 3.66<br>0.143 |
| Map resolution range (Å) | 2.8-7.0 |
| **Refinement** |  |
| Model resolution (Å)<br>FSC threshold | 3.7<br>0.5 |
| Map sharpening b-factor (Å<sup>2</sup>) | -186.3 |
| Model vs Map CC (mask) | 0.78 |
| Model composition<br>Non-hydrogen atoms<br>Protein residue<br>Water | 4162<br>545<br>3 |
| B factors (Å<sup>2</sup>)<br>Protein<br>Water | 32.23<br>16.76 |
| R.m.s. deviations<br>Bond lengths (Å)<br>Bond angles (°) | 0.003<br>0.564 |
| Validation<br>MolProbity score<br>Clashscore<br>Poor rotamers (%) | 1.80<br>8.9<br>0 |
| Ramachandran plot<br>Favored (%)<br>Allowed (%)<br>Disallowed (%) | 95.36<br>4.64<br>0 |

\*Values in parentheses indicate the pixel size in super-resolution.

**Table 2.**
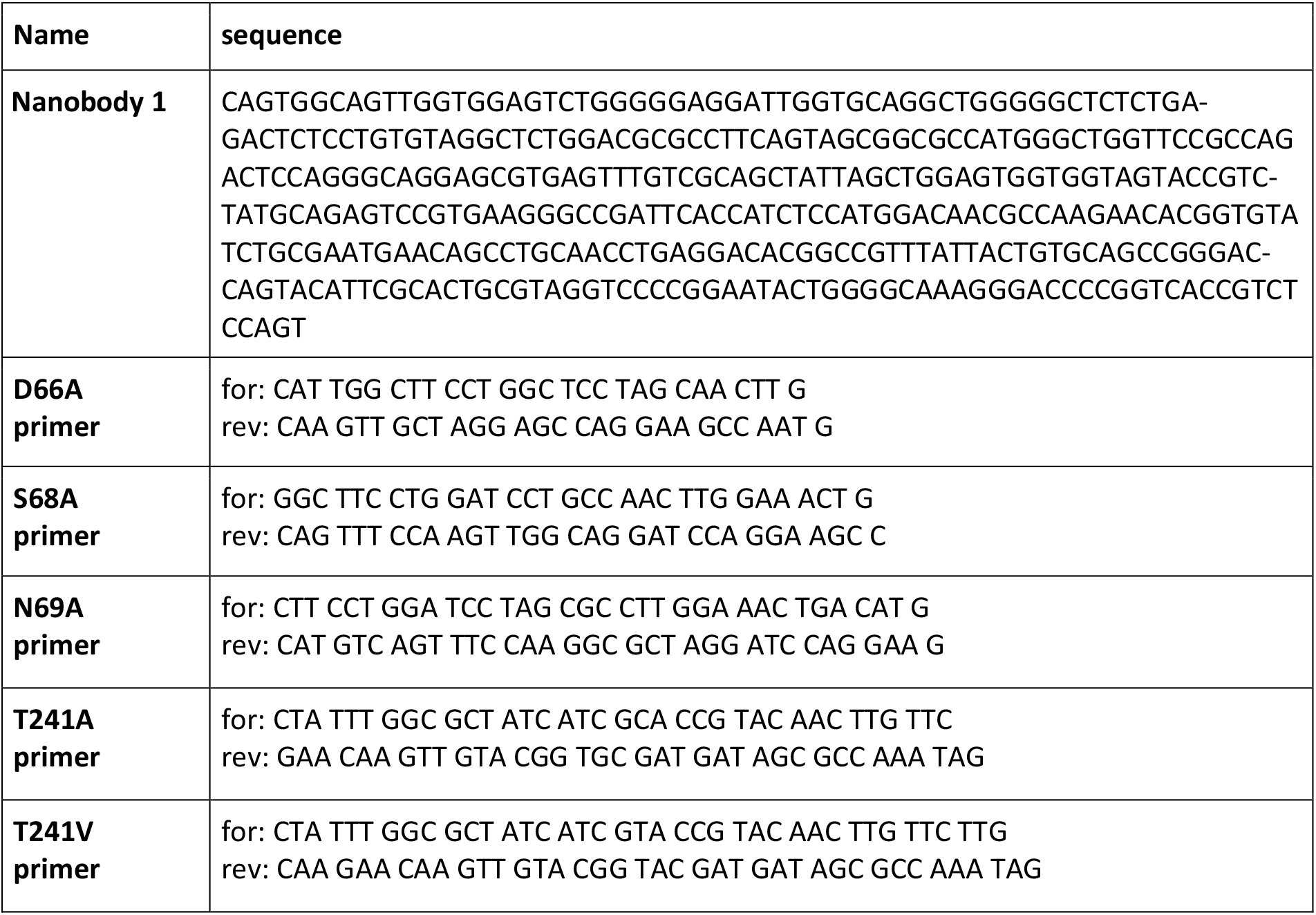
Nanobody and primer sequences.

### SiNRAT structure and presumable ion interactions

The SiNRAT structure in complex with Nb1 is depicted in Figure 2. It shows a protein sharing the conserved architecture of the SLC11 family with a core consisting of a pair of structurally related repeats of five transmembrane spanning segments that are oriented in the membrane with opposite directions in an arrangement that was initially observed in the amino acid transporter LeuT (Yamashita et al., 2005). In this architecture, the unwound centers of the first helix of each repeat together constitute a substrate binding site (Figure 2—figure supplement 4). The protein contains a total of twelve membrane-inserted helices akin to other eukaryotic family members and the prokaryotic EcoDMT (PDBID: 5M87) (Ehrnstorfer et al., 2017), which superimposes with an RMSD of 2.66 Å (Figure 2B, Figure 2—figure supplement 4). The nanobody binds to an extended interface located at the extracellular side. These interactions involve contacts of all three CDRs of the nanobody which bridge the core-domain consisting of α-helices 1-10 with α-helices 11-12 located at its periphery, thereby contacting the loops α1-2, α5-α6, α7-α8 and residues on α11 of SiNRAT and burying 1,682 Å^2^ of the combined molecular surface (Figure 2—figure supplement 4). Unlike in EcoDMT, in this complex α11 and α12 are detached from the core-domain on the extracellular part in a conformation resulting from the distinct state of the transporter that is presumably stabilized by Nb1 binding (Figure 2B, Figure 2—figure supplement 4).

The observed structure resembles an inward-occluded conformation of the classical SLC11 transporter DraNRAMP (PDBID: 8E60, DraNRAMP^occ^) where most of the structure retains a conformation observed in the inward-facing structure of the same protein (PDBID: 6D9W, DraNRAMP^inw^) except for a movement of the intracellular part of α-helix 1 (α1a), which has rearranged to close the access path to the ion binding site (Figure 3, Figure 3—figure supplement 1) (Bozzi et al., 2019b; Ray et al., 2022). This resemblance is reflected in an RMSD of 1.57 Å upon a superposition of SiNRAT with DraNRAMP^occ^ and 1.74 Å with DraNRAMP^inw^, whereas the RMSD of 2.71 Å compared to the outward-facing conformation of the same protein (PDBID: 6BU5, DraNRAMP^out^) is considerably larger (Figure 3B, Figure 3—figure supplement 1). As in DraNRAMP^occ^, the shorter α1a of SiNRAT is located closer to α6b compared to DraNRAMP^inw^, resulting in a narrowing of an aqueous cavity leading towards the supposed ion binding site, thereby separating this site from the intracellular environment (Figure 3C). We thus assume that the SiNRAT structure represents an intermediate between both endpoints of a transporter functioning by an alternate access mechanism that is closer to an inward-facing state. In its conformation, the binding site is sealed to both sides of the membrane by a thin gate composed of a single layer of residues *(i.e.* Asp 66, Asp 140, Thr 241, Tyr 243 and Thr 346) towards the inside and a thick gate (encompassing several layers of residues and spanning about 15 Å) towards the outside (Figure 3—figure supplement 1).

**Figure 3.**
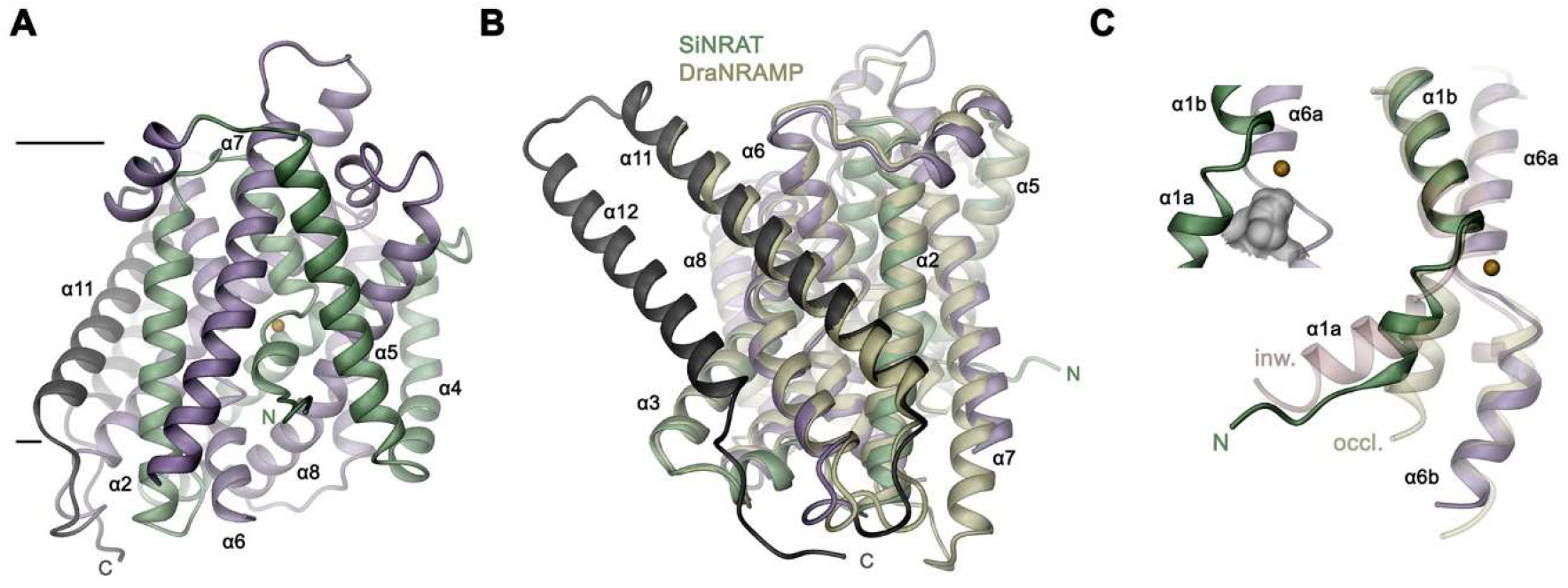
SiNRAT structure. (**A**) Ribbon representation of SiNRAT viewed from within the membrane with membrane boundaries indicated. (**B**) Superposition of SiNRAT with the protein DraNRAMP in an inward-occluded conformation. (**C**) Ion binding site in relation to the α-helices 1 and 6. The same helices of the superimposed structures of DraNRAMP^occ^ (PDBID: 6C3I, occl.) and DraNRAMP^inw^ (PDBID: 6D9W, inw.) are shown for comparison. Inset (left) shows the region surrounding the binding site with the molecular surface of the closed intracellular cavity displayed in grey. **A**, **C**, The position of the ion binding site is indicated by an orange sphere, **A-C**, The N and C-terminal repeats (α1-5 and a6-10) are colored in green and indigo respectively, the terminal helices α11 and 12 in dark grey. **Figure supplement 1.** Conformational properties of SLC11 transporters.

Although we have not explicitly added transported ions to our sample, we find residual cryo-EM density bridging the neighboring residues Asp 66 and Ser 68 which, as part of the supposed ion binding site, are both located on the unwound loop connecting α1a and b (Figure 4A). This density presumably either originates from water, the monovalent cation Na^+^ that is contained in the sample at a concentration of 200 mM, or traces of metal ions found in our solution. Due to its proximity to identified binding sites in structures of other family members, we use this position as reference to analyze plausible protein substrate interactions. In this putative site, Asp 66 is particularly striking, since it is universally conserved and was shown to constitute a key component for ion coordination in the entire SLC11 family. In contrast, Ser 68 is unique to NRATs and substituted by a Gly in all other members. The comparison of the SiNRAT site with the ion interaction in DraNRAMP^occ^ shows common features but also differences that may underlie the altered substrate preference of SiNRAT. In both structures, the water-filled paths to the metal ion binding site are sealed from both sides of the membrane and bound ions would thus be segregated from their aqueous environment, although the occlusion towards the intracellular solution is more pronounced in DraNRAMP^occ^ (Figure 3C and 4B, C, Figure 3—figure supplement 1). In its binding site, the interacting transition metal ion in DraNRAMP^occ^ is enclosed by protein residues except for a small water-filled pocket contacting the bound ion on one side (Ray et al., 2022). The sidechains of Asp 56 and Asn 59 on α1 in conjunction with the backbone carbonyls of Ala53 and Ala 227 tightly surround the bound Mn^2+^. Additionally, the thioether group of Met 230 contributes a soft ligand for ion interactions (Figure 4C). Finally, a water-mediated interaction with the backbone carbonyl of Gln 378 on α10 completes the octahedral coordination of the ion (Figure 4C). In contrast, the equivalent region in SiNRAT contains an elongated, approximately cylindrical pocket of predominant polar character that is about 10 Å long and 4-5 Å wide and that encloses a three-times larger volume than in DraNRAMP (Figure 4B). This pocket appears of sufficient size to mediate metal ion interactions with protein residues and surrounding water molecules. Despite the inability to pinpoint the exact location of bound ions, we find residues of the consensus binding site to line about half of this pocket (Figure 4B). These include the previously mentioned Asp 66 and Ser 68, both bordering residual density in our cryo-EM data and the conserved Asn 69, all located on α1. On α6, the interacting residues include the conserved Ala 238, whose backbone carbonyl points towards the pocket and Thr 241, which replaces the conserved methionine in NRAMP transporters and whose sidechain contributes a hard ligand for ion interactions (Figure 4B). Distinct from DraNRAMP, the glutamine on α10 involved in ion binding is replaced by a serine (Ser 403), whose shorter side chain does not contact the aqueous pocket and thus would not be available for metal ion interactions. Instead, we find the acidic sidechain of Asp 140 located on α3 to line the cavity close to the predicted ion binding site at a position that contains a threonine (Thr 130) in DraNRAMP. This acidic sidechain, which is conserved in NRATs, would increase the negative charge density of the region by offering favorable coulombic interactions that stabilize the additional positive charge of Al^3+^ (Figure 4B, Figure 3—figure supplement 1). Thus, besides the replacement of the conserved methionine of the transition metal ion binding site, we find an expanded interaction region with increased polar character and negative charge density, which together may account for the altered substrate preference and stabilize trivalent cations.

**Figure 4.**
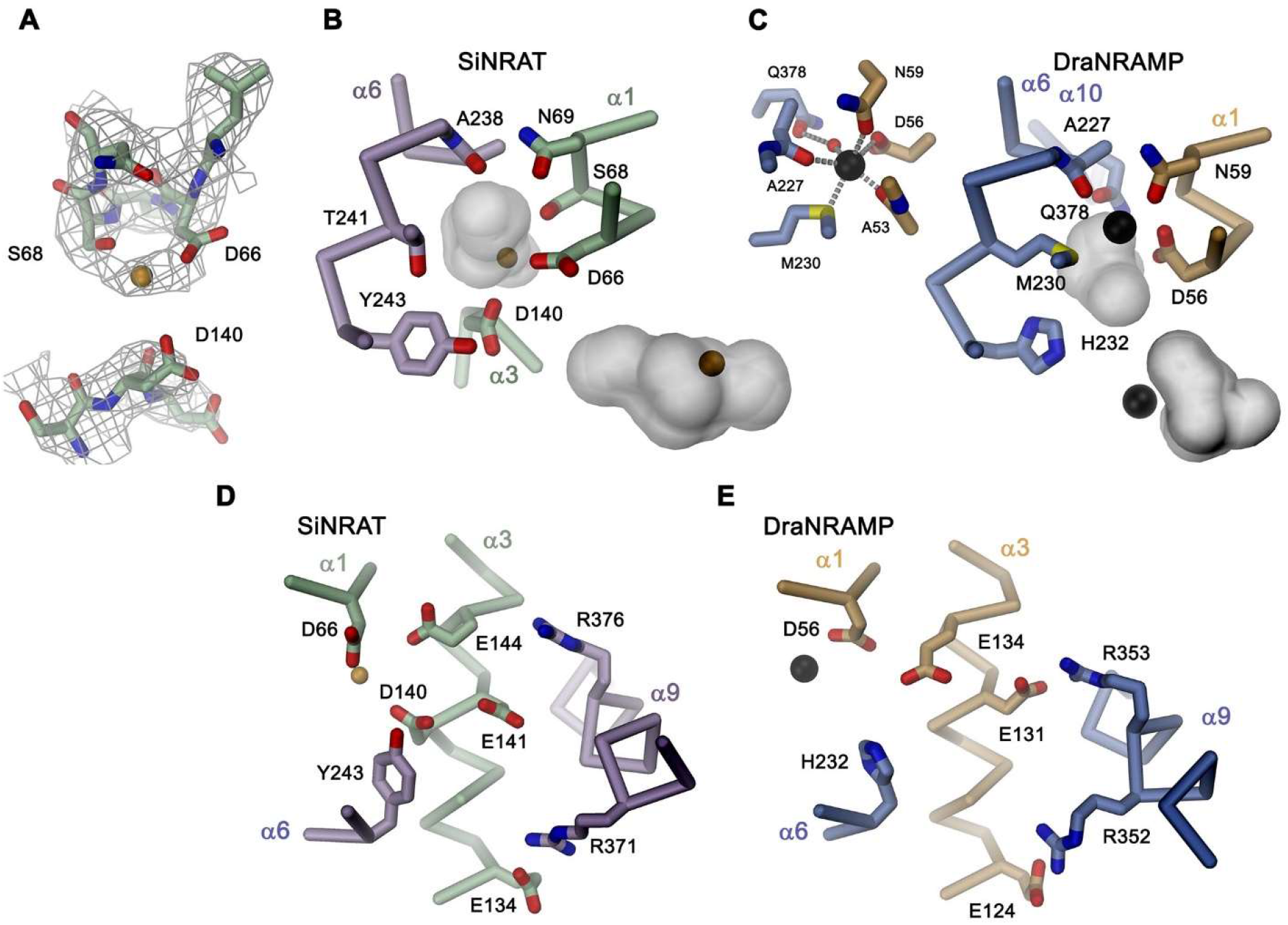
SiNRAT metal ion binding site. (**A**) Selected parts of the SiNRAT ion binding region with cryo-EM density superimposed. Top, loop connecting helices α1a and b. Residual density between Asp 66 and Ser 68 was used to define the presumable position of the ion binding site (with a bound ion represented as orange sphere). Bottom, density around Asp 140 located on α3. Comparison of the metal ion binding site in SiNRAT (**B**) and DraNRAMP^occ^ (PDBID: 8E60) (**C**) viewed from within the membrane. Shown are the presumed positions of bound metal ions (spheres) and the surrounding protein as Cα trace with selected interacting side and main chain atoms labeled and displayed as sticks. In both cases the occluded cavity surrounding the bound ion that is presumably filled by water is shown as grey surface. Insets (bottom right), show the same cavity viewed from the cytoplasm. In **C**, a second inset (top right) shows the coordination geometry of the Mn^2+^ ion bound to DraNRAMP. **D**, **E** Comparison of the corresponding regions bearing residues that were assigned a potential relevance for proton transport in NRAMPs in the uncoupled SiNRAT (**D**) and the H^+^-coupled DraNRAMP (**E**). The major difference concerns a conserved His on α6 (His 232 in DraNRAMP) that is located intracellular to the metal ion binding site, that is a Tyr in SiNRAT (Tyr 243). **A**-**E**, Residues of the N-and C-terminal repeats of both proteins are shown in unique colors.

Besides the altered ion selectivity, the second feature that distinguishes SiNRAT from classical NRAMPs is the lack of proton coupling, which is also shared by prokaryotic NRMTs of the same family (Ramanadane et al., 2022). In NRAMPs, proton transport was assigned to acidic and basic residues located on the α-helices 1, 3, 6, and 9, most of which are replaced by hydrophobic sidechains in NRMTs (Bozzi et al., 2019a; Bozzi et al., 2020; Ehrnstorfer et al., 2017; Ramanadane et al., 2022). In contrast, many of these residues are conserved in NRATs, which are evolutionally closer to NRAMPs (Figure 4D, E). This is the case for acidic residues located on α3 *(i.e.* Glu 134, Glu 141 and Glu 144), where we even find an additional acidic residue lining the ion binding site (i.e. Asp 140). Two residues on α9 (Arg 371 and Arg 376), although not located at equivalent positions, form corresponding interactions as found in the H^+^-coupled DraNRAMP. A pronounced difference between NRATs and NRAMPs concerns the substitution of a conserved histidine (His 232 in DraNRAMP) (Ehrnstorfer et al., 2017; Lam-Yuk-Tseung et al., 2003) on α6b with a tyrosine (Tyr 243), which could contribute to the altered transport phenotype although we expect also other parts of the protein to play a role in this process.

### Functional properties of SiNRAT binding site mutants

After defining the structural features of the putative metal ion binding site of SiNRAT, we decided to study the effect of point mutants on the transport properties of the protein. To this end, we have put our focus on residues of the consensus binding site located on α-helices 1 and 6, whose equivalent positions were found to contribute to ion interaction in transition metal ion transporters of the SLC11 family (Figure 5A). All investigated mutants were purified and either reconstituted into liposomes for uptake studies of Mn^2+^ or used for ITC experiments in their detergent-solubilized form. Upon mutation of the residues Asp 66 or Asn 69 to alanine, we found a severely compromised transport of Mn^2+^, consistent with equivalent results obtained for other pro- and eukaryotic transition metal ion transporters (Figure 5B, C) (Bozzi et al., 2019b; Ehrnstorfer et al., 2014; Ehrnstorfer et al., 2017). We next turned our attention towards Thr 241, which has substituted the conserved methionine found in transition metal ion transporters and provides a hard ligand for metal ion interactions. In this case, we have mutated the residue to alanine to truncate the sidechain and valine to preserve its volume but remove its polar character. In both cases, we found a similar strong negative impact of the respective mutations on Mn^2+^ transport (Figure 5D, E). For this residue, we were also interested whether a mutation would affect interactions with trivalent cations and thus investigated binding of Al^3+^ to the mutant T241V by ITC. Unlike WT, we did not detect any specific heat exchange, which emphasizes the importance of the residue for Al^3+^ interactions (Figure 5F). The absence of a detectable signal in T241V, also underlines the correspondence of the WT data to Al^3+^ binding to the consensus site (Figure 1H). Finally, we have studied a mutation of Ser 68 to alanine, which concerns a residue that is unique to NRATs and which was found in the SiNRAT structure to contribute to potential ion interactions (Figure 4A). Unlike previous mutations, we did in this case not observe an obvious phenotype on Mn^2+^ transport, which proceeds with similar kinetics as WT (Figure 5G, H) and whose transport is competed by the addition of Mg^2+^ and Ca^2+^ in a similar manner (Figure 5I, J). Unlike WT, we did not detect a competition by Ga^3+^ in equivalent experiments (Figure 5K). We thus set out to investigate the binding of Al^3+^ by ITC but did not find any indication for its interaction with the mutant (Figure 5L). Taken together, our experiments on the SiNRAT mutant S68A showed distinct phenotypes for the interaction with di- and trivalent cations, which points towards structural differences in their coordination with more stringent requirements for Ga^3+^ and Al^3+^ binding, latter being the presumable substrate under physiological conditions.

**Figure 5.**
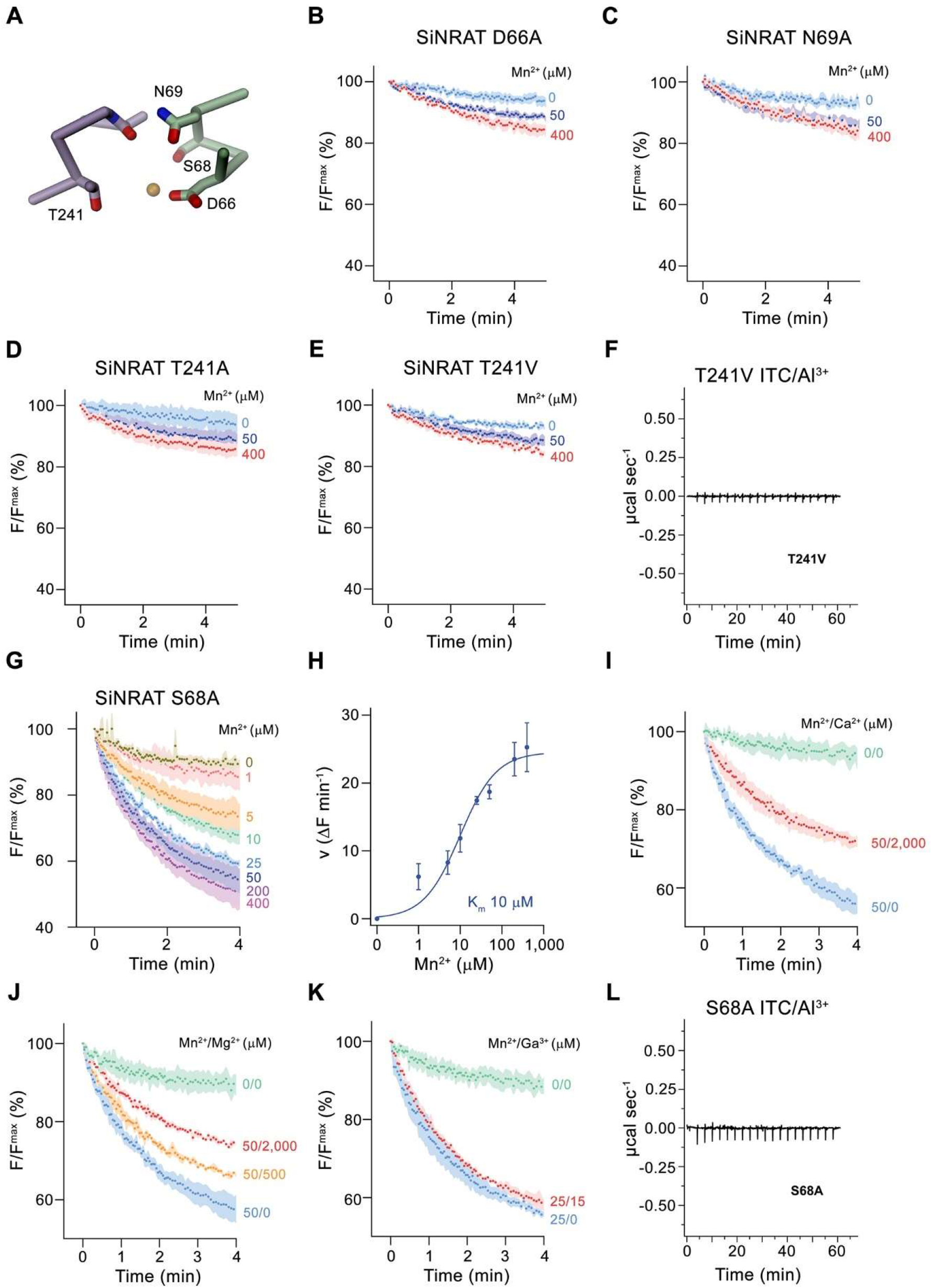
Functional properties of binding site mutants. (**A**) Cα trace of the metal ion binding region of SiNRAT with selected residues involved in ion coordination shown as sticks. **B-E**, Mn^2+^ transport into proteoliposomes of SiNRAT metal ion binding site mutants. (**B**) D66A (4 experiments from 3 independent reconstitutions for all conditions); (**C**) N69A (3 experiments from 3 independent reconstitutions); (**D**) T241A (4 experiments from 3 independent reconstitutions for all conditions); (**E**) T241V (3 experiments from 3 independent reconstitutions for all conditions). (**F)** Thermogram of Al^3+^ titrated to the SiNRAT mutant T241V obtained from isothermal titration calorimetry experiments. (**G**) Mn^2+^ transport into proteoliposomes containing the SiNRAT mutants S68A (3 experiments from 2 independent reconstitutions for all conditions). (**H**) Mn^2+^ concentration dependence of transport by the mutant S68A. Data show mean of initial velocities derived from individual traces of experiments displayed in **G**, errors are s.e.m., the solid line shows the fit to a Michaelis–Menten equation with an apparent *K*_m_ of 10 μM. **I-K,** Mn^2+^ transport in presence of other multivalent cations show interaction of the mutated binding site with Ca^2+^ and Mg^2+^ but not with Ga^3+^ at indicated ion concentrations. (**I**) Ca^2+^ (3 experiments from 2 independent reconstitutions for all conditions); (**J**) Mg^2+^ (3 experiments from 2 independent reconstitutions); (**K**) Ga^3+^ (3 experiments from 2 independent reconstitutions). (**L)** Thermogram of Al^3+^ titrated to the SiNRAT mutant S68A obtained from isothermal titration calorimetry experiments. **B-E**, **G**, **IK**, Uptake of Mn^2+^ was assayed by the quenching of the fluorophore calcein trapped inside the vesicles. Panels show mean of indicated number of experiments, errors are s.e.m.. Fluorescence is normalized to the value after addition of substrate (t=0). Applied ion concentrations are indicated.

## Discussion

Plants growing on acidic soils have to cope with elevated concentrations of Al^3+^ which, as strong Lewis acid, forms tight interactions with proteins and phospholipids thereby compromising membrane integrity (Chauhan et al., 2021). As one of the mechanisms to combat Al^3+^ toxicity, certain plants have evolved a branch of the NRAMP family with altered substrate preference termed NRATs, which are capable of transporting the trivalent cation (Xia et al., 2010). In our study, we have determined the structure and transport properties of the NRAT of *Setaria italica* termed SiNRAT. Using fluorescence-based *in vitro* transport assays from proteoliposomes containing the purified transporter, our study has characterized SiNRAT as a protein with broad substrate specificity. We have shown transport of different divalent cations such as Mn^2+^ Ca^2+^ and Mg^2+^ with micromolar K_M_ (Figure 1A-F). Although the direct assay of transport of trivalent ions such as Ga^3+^ and Al^3+^ was prohibited by technical limitations, we have demonstrated the interaction of SiNRAT with both ions, which defines them as plausible substrates (Figure 1G-I). The broad substrate selectivity is consistent with previous experiments on the metal ion transporters EcoDMT and DraNRAMP (Bozzi et al., 2016; Ehrnstorfer et al., 2017) and the Mg^2+^ transporter EleNRMT (Ramanadane et al., 2022). Attempts in these proteins to change their transport properties by mutagenesis, while deteriorating selectivity, have usually retained the capability of mutants to permeate Mn^2+^ (Bozzi et al., 2016; Ramanadane et al., 2022), whose transport appears to have lower structural requirements compared to other divalent cations. By the placement of a methionine contributing a soft ligand for transition metal ion interaction, the substrate binding site of classical NRAMPs has evolved to prevent interactions with alkaline earth metal ions (Bozzi et al., 2016) (Figure 6A). This is illustrated in the mutation of this residue to alanine, which has turned Ca^2+^ into a substrate without strongly affecting Mn^2+^ transport (Bozzi et al., 2016; Ramanadane et al., 2022). Similarly, NRMTs transport the small alkaline earth metal ion Mg^2+^ by increasing the volume of their binding site to interact with the hydrated cation, without compromising Mn^2+^ transport (Ramanadane et al., 2022) (Figure 6B). A comparable feature is observed for SiNRAT, where the protein appears to have evolved to transport trivalent cations without imposing a strong selection against divalent metal ions (Figure 1). The SiNRAT structure has shed light into the basis for the altered selectivity of a transporter, whose evolutionary distance to classical NRAMPs is much smaller than to NRMTs (Lu et al., 2018; Ramanadane et al., 2022). Although our structure did not contain trivalent ions, we assigned their binding site based on the structural equivalence to the transition metal ion transporters DraNRAMP and ScaDMT and the Mg^2+^ transporter EleNRMT (Ehrnstorfer et al., 2014; Ramanadane et al., 2022; Ray et al., 2022). Residual density at the supposed location in the SiNRAT map provides further evidence for this assignment. Similar to NRMTs, and distinct from classical NRAMPs, the ion binding site of SiNRAT is embedded in a larger aqueous cavity, which offers direct interactions with protein residues and others that are mediated via hydrating waters (Figure 4B, C, Fig. 6C) (Ramanadane et al., 2022). Unlike the inward-facing structure of EleNRMT, this cavity is sealed from both sides of the membrane (Figures 2C and 3B). Within the canonical binding site of SiNRAT located on α1 and α6, several residues have been replaced to provide additional interactions and the presence of an extra acidic residue on α3 in interaction distance has increased the negative charge density to compensate for the higher net charge of the transported ion (Figure 4B). In light of the described features, a proposed conversion of a classical plant NRAMP facilitating the transport of transition metals by the mutation of the consensus binding motif on α1 and α6 is presumably insufficient to convert the protein into an efficient Al^3+^ transporter (Lu et al., 2018). A related attempt to convert Mg^2+^ into a transported substrate of the prokaryotic metal ion transporter EcoDMT, by introducing corresponding mutations found in NRMTs was unsuccessful (Ramanadane et al., 2022), which emphasizes the relevance of residues outside a narrow signature region to confer substrate selectivity.

**Figure 6.**
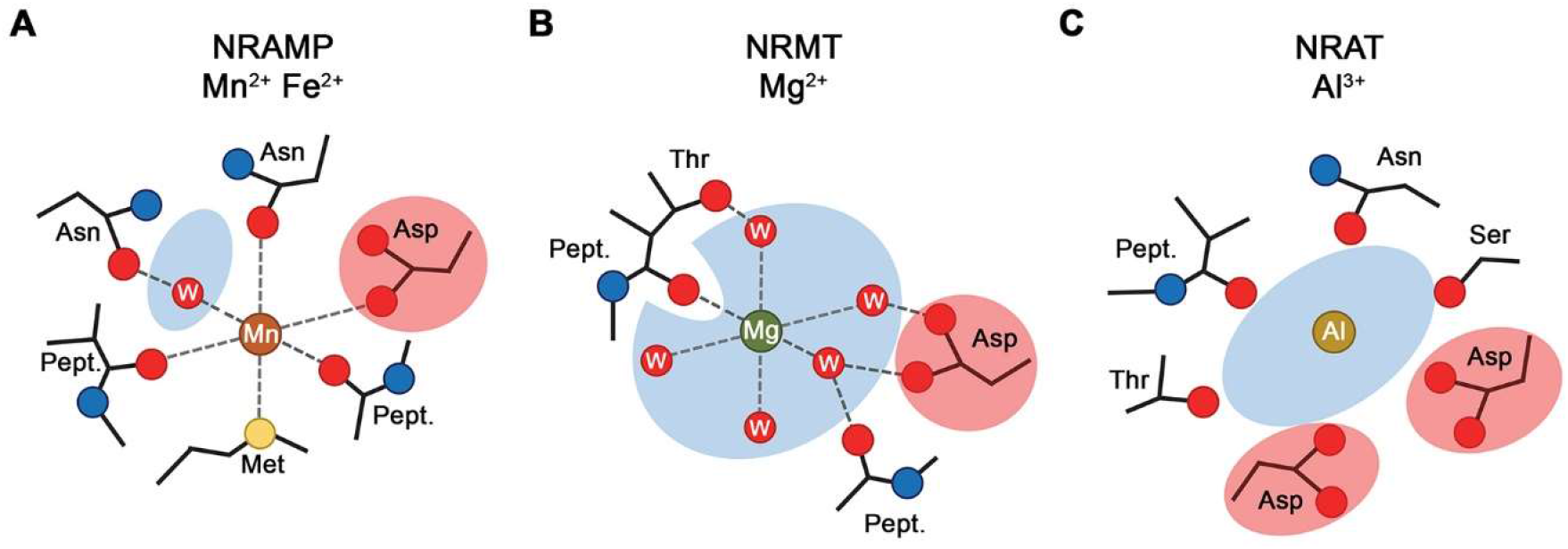
**S**electivity and metal ion coordination in different clades of the SLC11 family. (**A**) Coordination of Mn^2+^ in transition metal ion transporters (NRAMPs) of the SLC11 family as defined in the occluded conformation of the prokaryotic DraNRAMP (PDBID: 8E60). The octahedral coordination geometry is well-defined. The largely dehydrated metal ion is predominantly forming direct protein interactions except for one interaction that is mediated by a bound water molecule. The thioether of a methionine serves as soft ligand in transition-metal ion coordination. (**B**) Putative Mg^2+^ interaction in an NRMT. The interactions were obtained from the inward-facing conformation of the protein EleNRMT (PDBID: 7QIA). The location of the metal ion was defined by bound Mn^2+^ obtained from anomalous X-ray scattering experiments, the position of coordinating water molecules are modeled. Owing to the larger volume of the binding site, the ion has retained most of its first coordination shell and the majority of ion-protein interactions are presumably indirect. (**C**) Putative coordination of Al^3+^ in the transporter SiNRAT. The detailed binding position is not defined with confidence but was inferred from residual density in the SiNRAT cryo-EM map. The ion is placed in an aqueous cavity and likely undergoes interactions with the protein that are either direct or mediated by water molecules. A second presumably deprotonated aspartate increases the negative charge density in the binding site and thus stabilizes the greater charge density of the trivalent metal ion. **A-C**, Interacting waters (W) are labeled, an aqueous cavity is indicated in blue, negatively charged groups involved in metal ion coordination are highlighted in red. Pept. refers to the interaction with the carbonyl of a peptide bond.

As in EleNRMT, transport by SiNRAT appears not to be coupled to protons, although, unlike former, many of the residues that were proposed to contribute to H^+^ coupling in NRAMPs are preserved in the protein (Bozzi et al., 2019a; Bozzi et al., 2020; Ehrnstorfer et al., 2017). A prominent exception concerns the replacement of a conserved His on α6 located in proximity to the ion binding site by a Tyr (i.e. Tyr 243), whose hydroxyl group also lines the cavity. Remarkably, the mutation of the equivalent position in EcoDMT has abolished H^+^ transport (Ehrnstorfer et al., 2017). The observed broad substrate specificity of SiNRAT contrasts a previous study on the prototypic transporter NRAT1 from rice which, upon overexpression in yeast, was proposed to selectively transport Al^3+^ but not the transition metal ion Mn^2+^ (Xia et al., 2010). This apparently contradicting observation could reflect the distinct ways of how transport was assayed in both studies. We thus remain cautious with the assignment of substrates transported by NRATs in a physiological context as the protein might not be efficient in concentrating Mn^2+^ inside cells, potentially as a consequence of the absent coupling to an energy source. Still, in light of our data it is unclear how the presence of other divalent ions would not interfere with Al^3+^ transport in a physiological context. In summary our study has provided the first structural insight into the transport of trivalent cations and it revealed the architecture of a protein that is part of the defense against Al^3+^ toxicity in plants, which continues to be a large challenge for farming on acidic soil. Since it was previously shown that the increased expression of proteins involved in the regulation of aluminum toxicity can lead to better aluminum tolerance (Maron et al., 2013) it will be interesting to investigate whether the overexpression of NRATs would increase the tolerance of plants to elevated Al^3+^ levels.

## Materials and Methods

### Key resources table

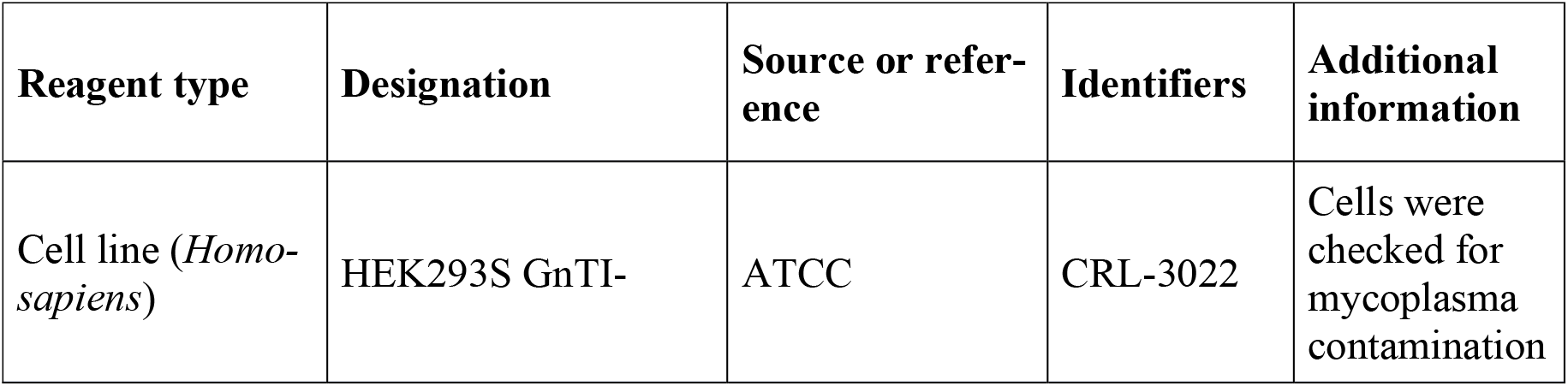
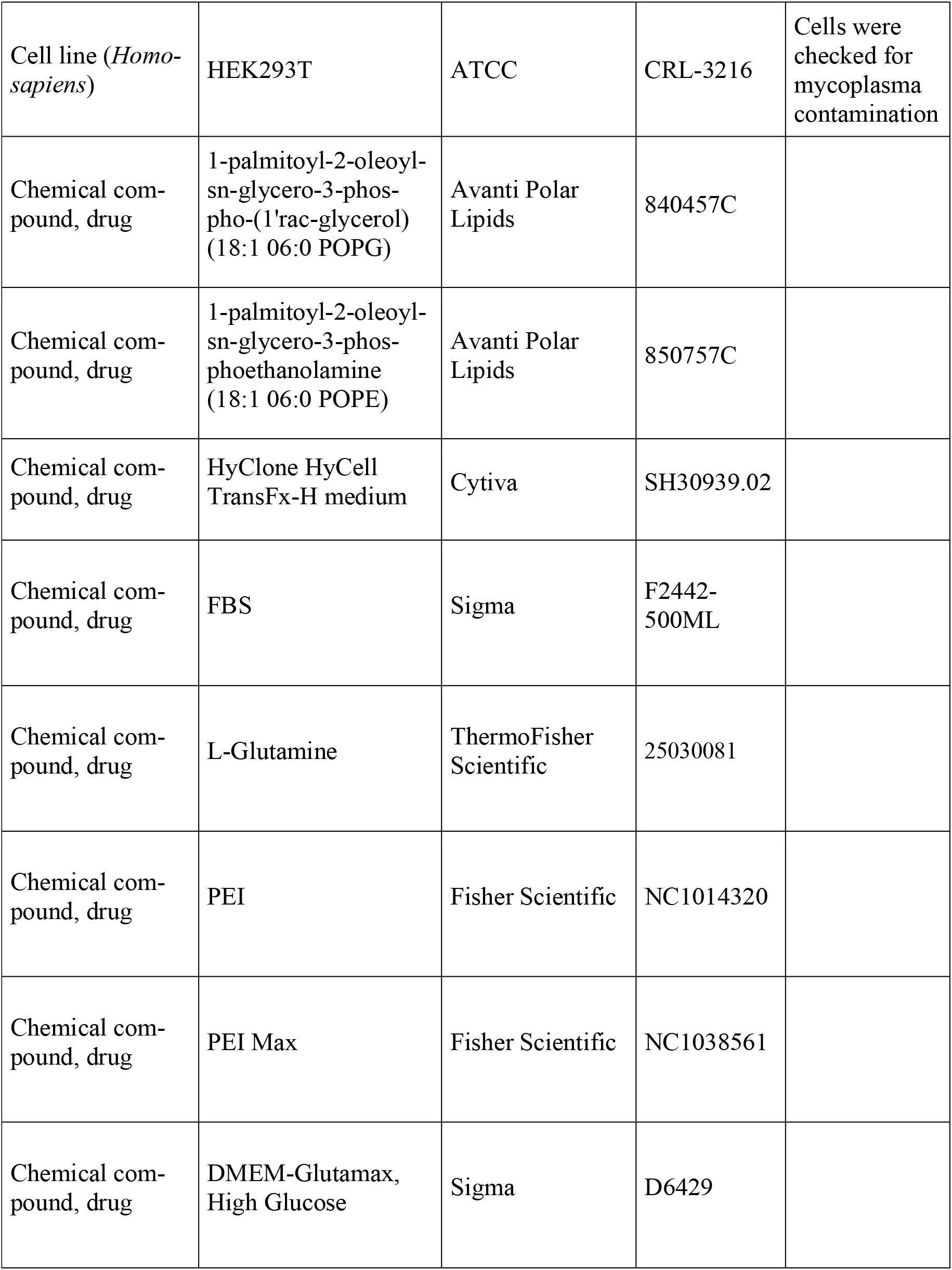
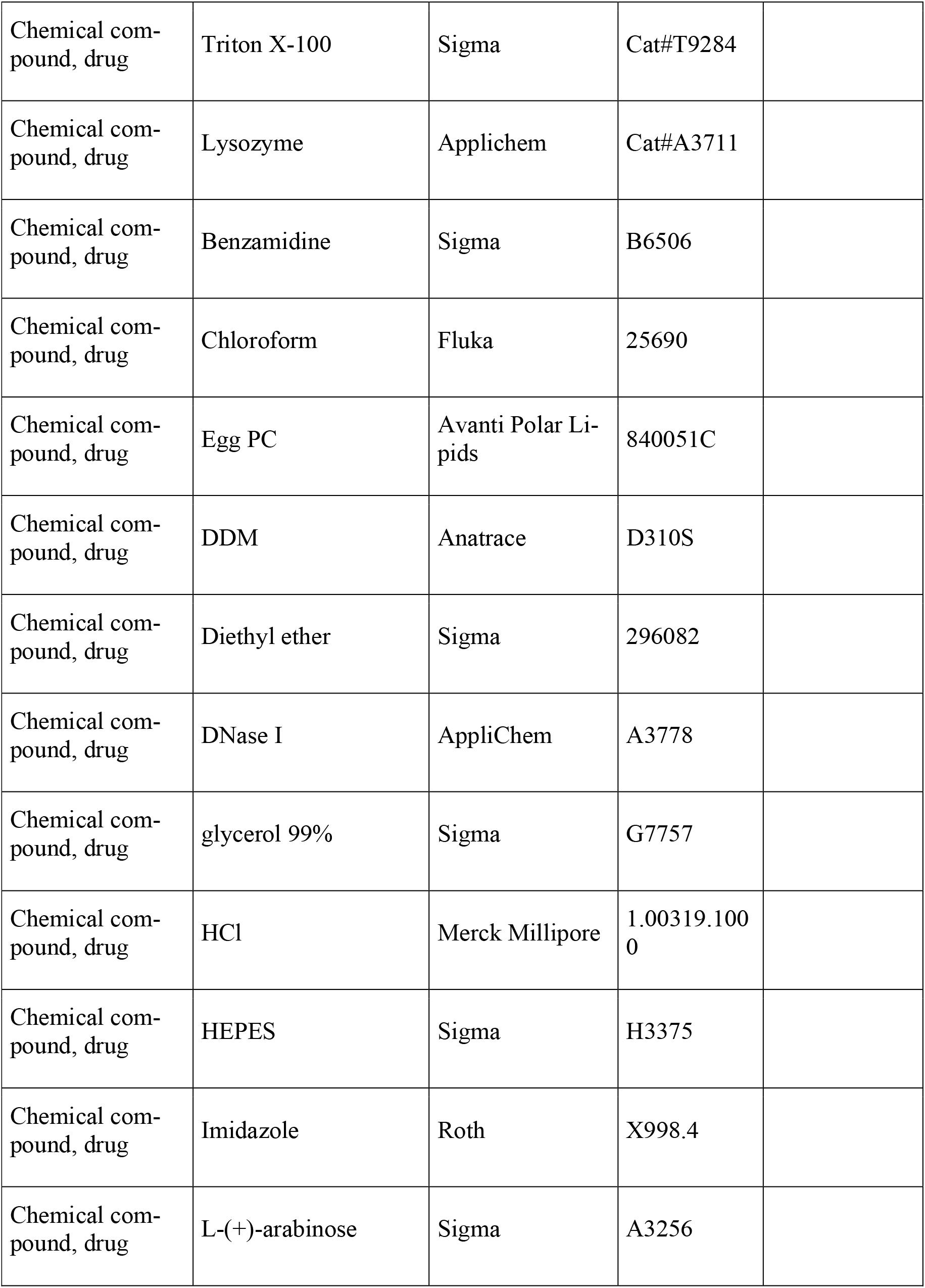
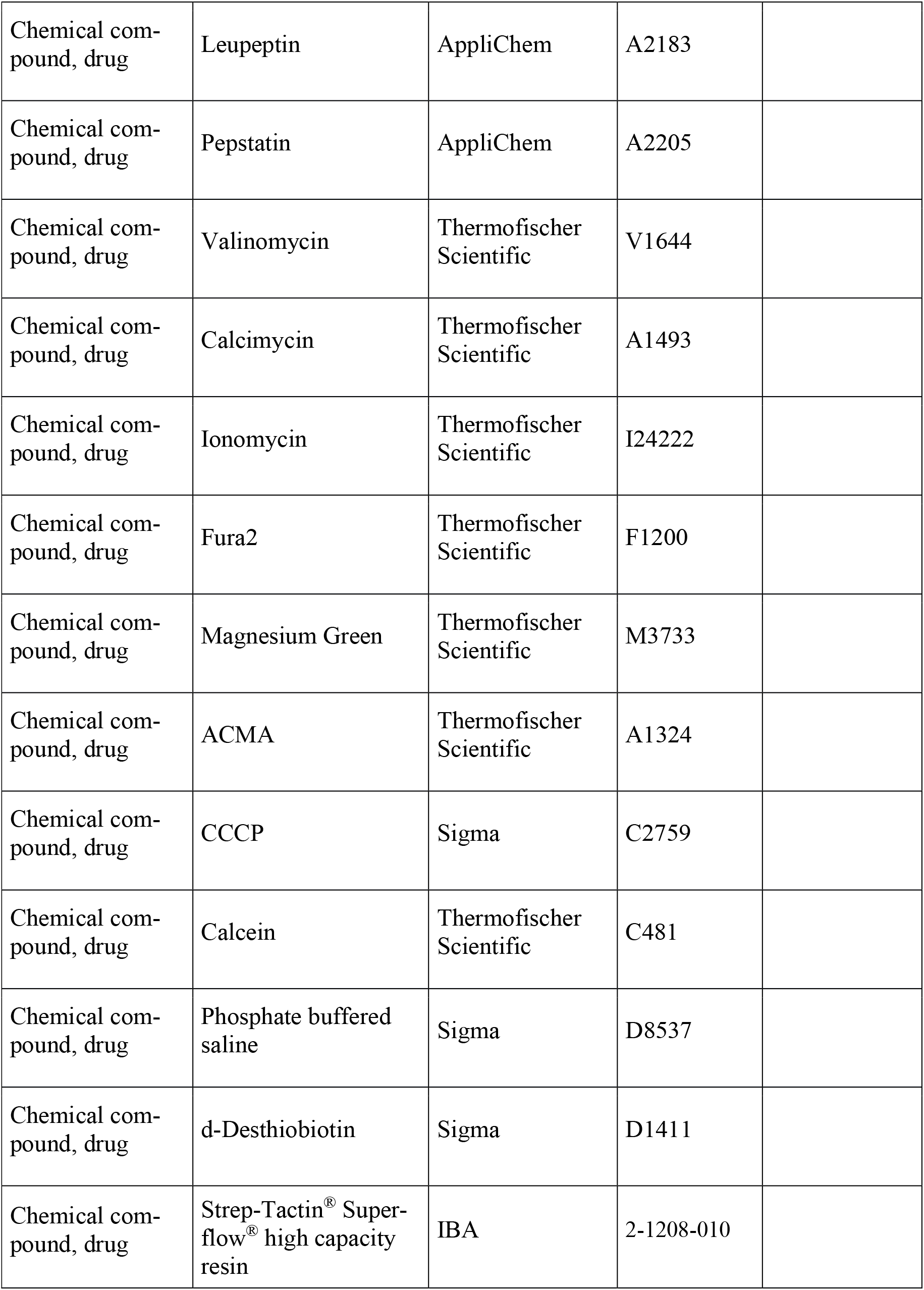
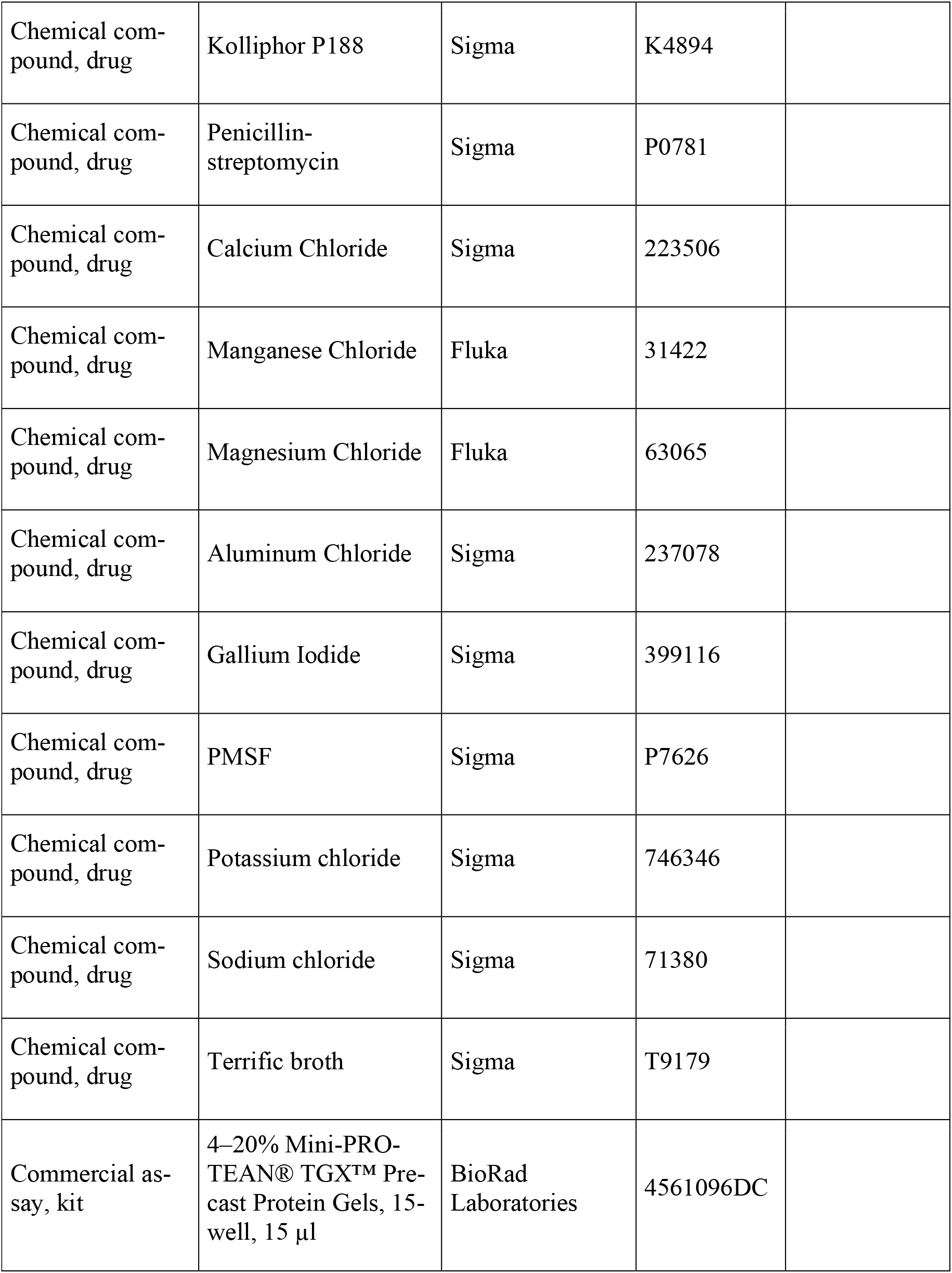
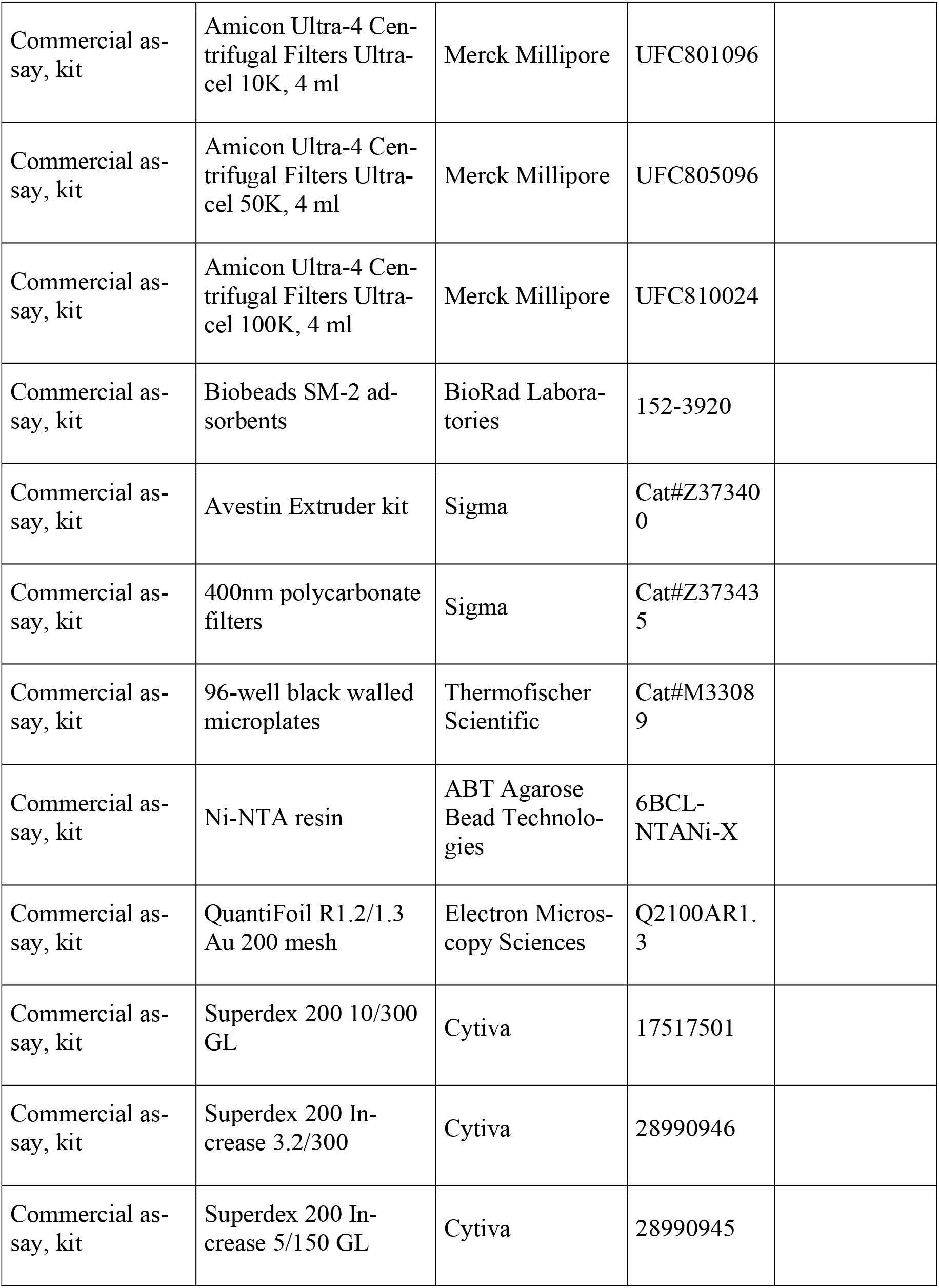
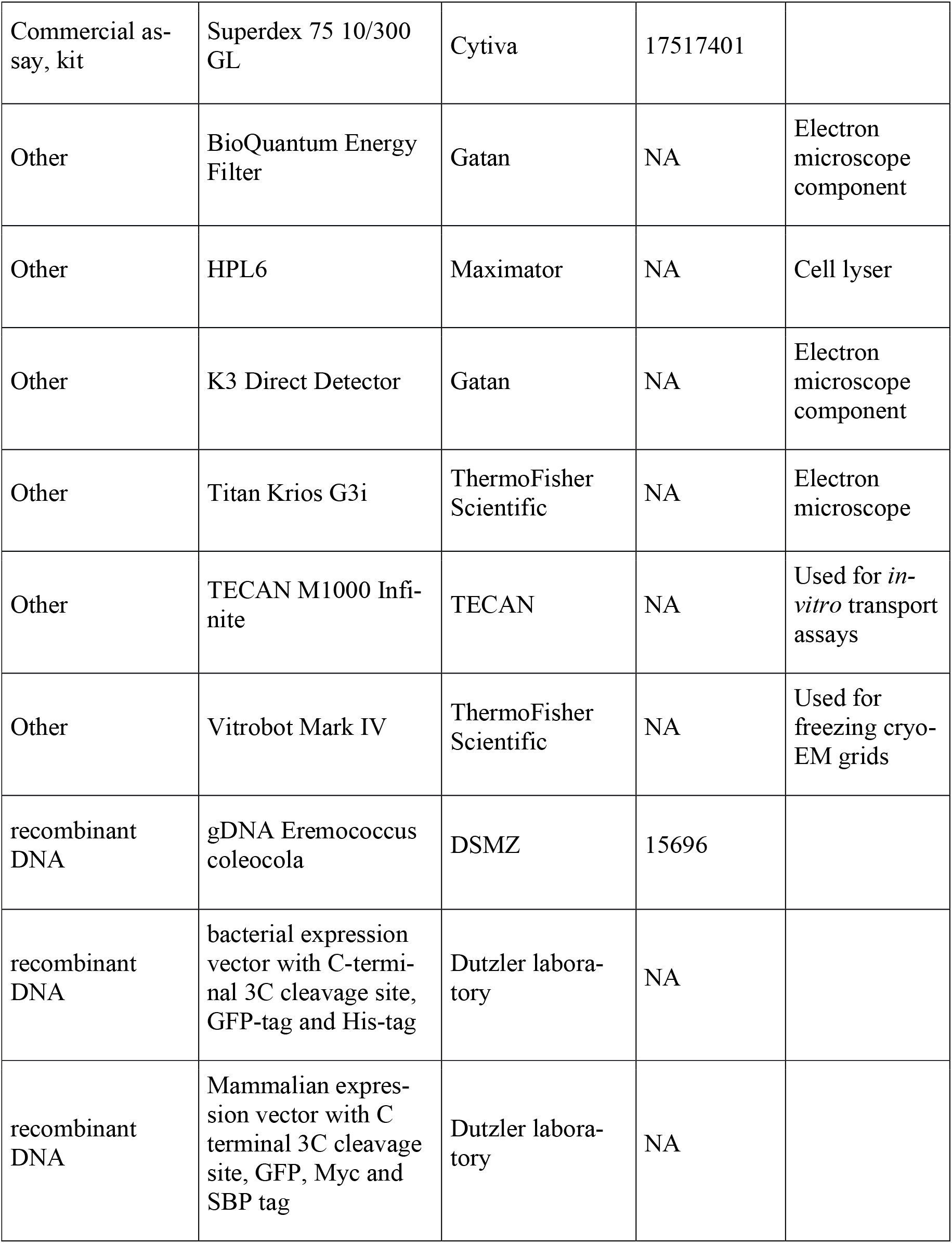
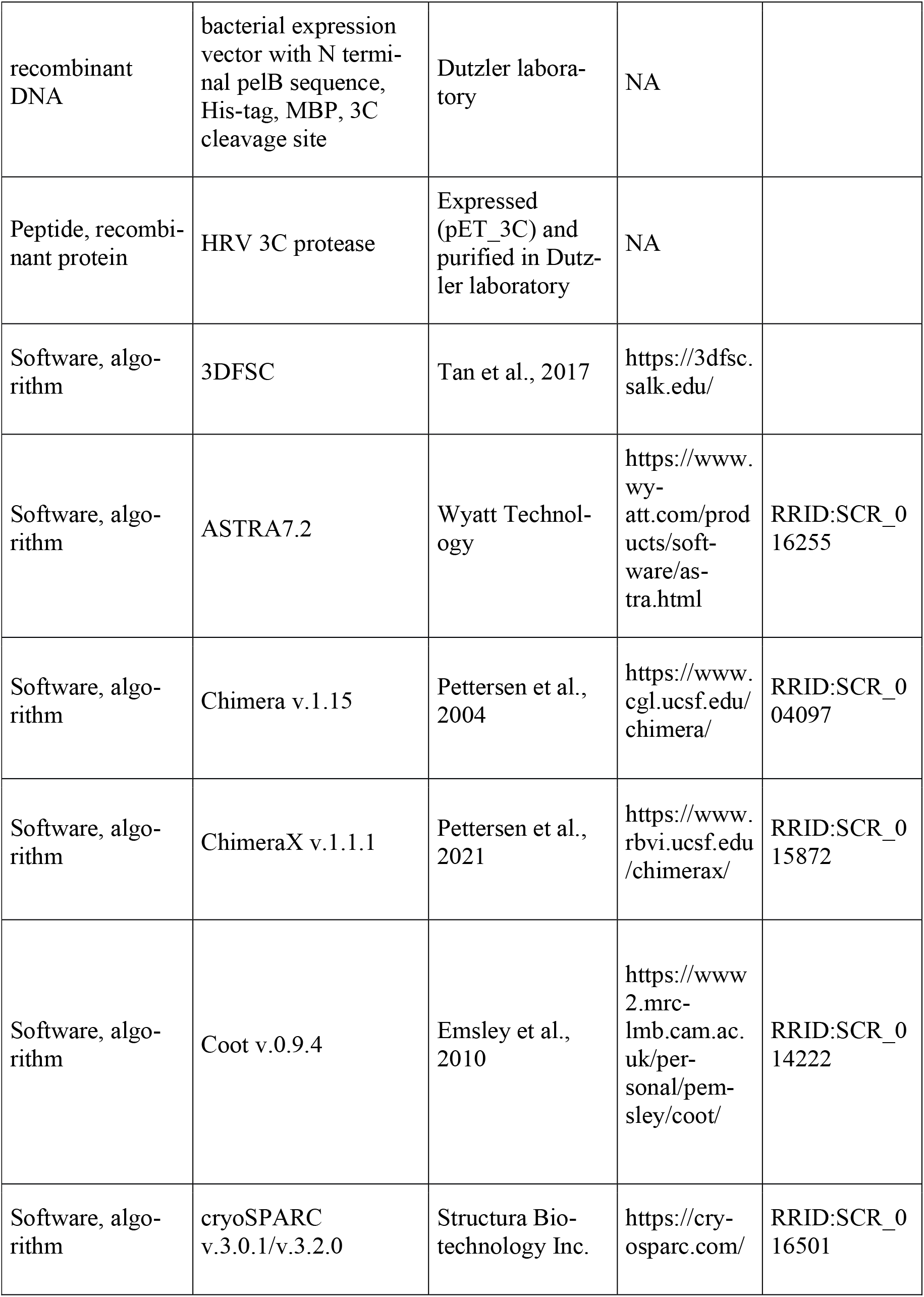
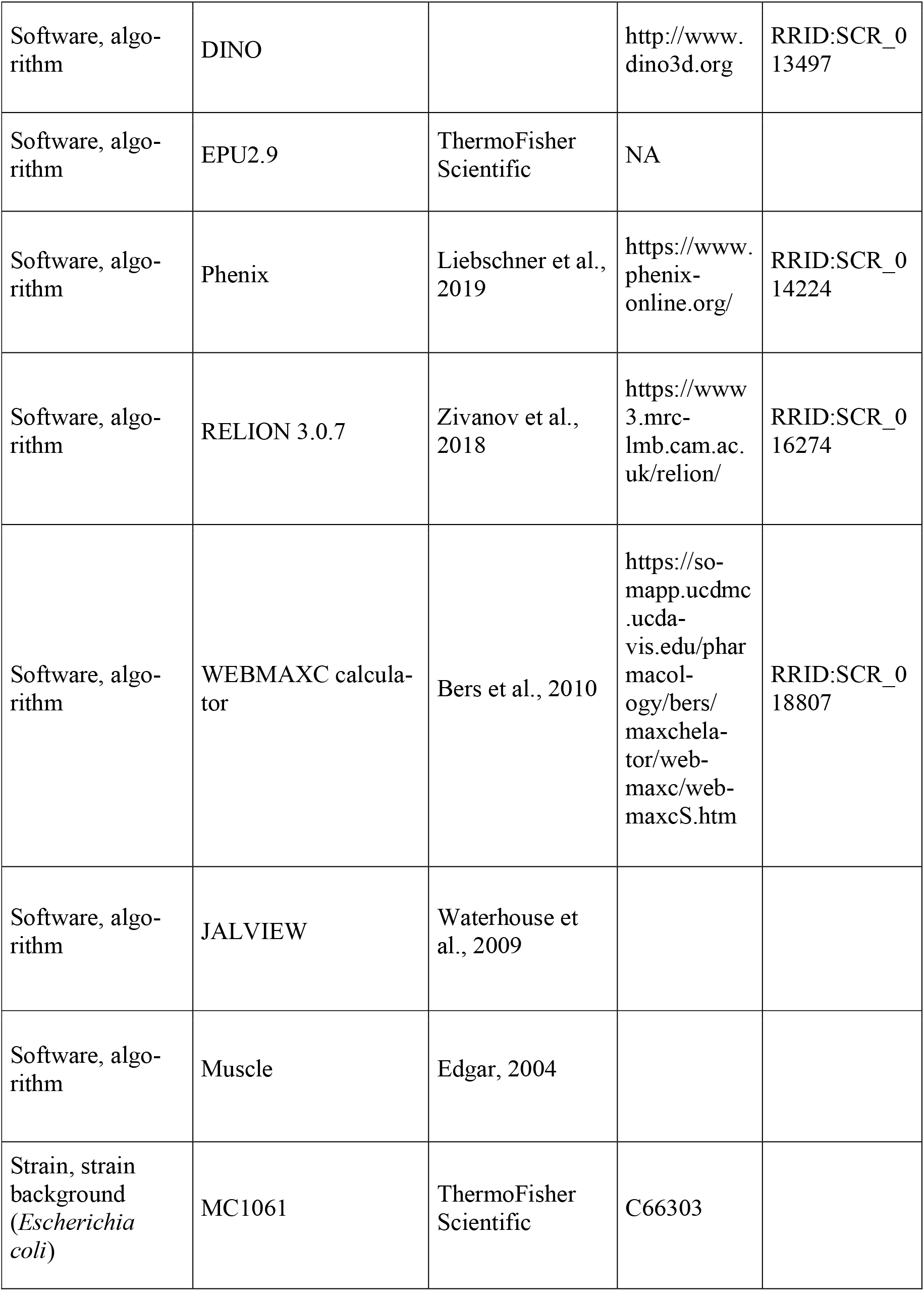

### Expression and purification of EcoDMT

EcoDMT and its mutants were expressed and purified as described previously (Ehrnstorfer et al., 2017). Briefly, the vector EcoDMT-pBXC3GH was transformed into *E. coli* MC1061 cells and a preculture was grown in TB-Gly medium overnight using ampicillin (100 μg/ml) as a selection marker for all expression cultures. The preculture was used at a 1:100 volume ratio for inoculation of TB-Gly medium. Cells were grown at 37 °C to an OD600 of 0.7-0.9, after which the temperature was reduced to 25 °C. Protein expression was induced by addition of arabinose to a final concentration of 0.004% (w/v) for 12-14 h at 18 °C. Cells were harvested by centrifugation for 20 min at 5,000 g, resuspended in buffer RES (200 mM NaCl, 50 mM KPi pH 7.5, 2 mM MgCl_2_, 40 μg/ml DNAseI, 10 μg/ml lysozyme, 1μM leupeptin, 1 μM pepstatin, 0.3 μM aprotinin, 1mM benzamidine, 1mM PMSF) and lysed using a high pressure cell lyser (Maximator HPL6). After low speed centrifugation (10,000 g for 20 min), the supernatant was subjected to a second centrifugation step (200,000 g for 45 min). The pelleted membrane was resuspended in buffer EXT (200 mM NaCl, 50 mM KPi pH 7.5, 10% glycerol) at a concentration of 0.5 g of vesicles per ml of buffer. The purification of EcoDMT and its mutants was carried out at 4 °C. Isolated membrane fractions were diluted 1:2 in buffer EXT supplemented with protease inhibitors (Roche cOmplete EDTA-free) and 1% (w/v) n-decyl-β-D-maltopyranoside (DM, Anatrace). The extraction was carried out under gentle agitation for 2 h at 4 °C. The lysate was cleared by centrifugation at 200,000 g for 30 min. The supernatant supplemented with 15 mM imidazole at pH 7.5 was subsequently loaded onto NiNTA resin and incubated for at least 1 h under gentle agitation. The resin was washed with 20 column volumes (CV) of buffer W (200 mM NaCl, 20 mM HEPES pH 7, 10% Glycerol, 50 mM imidazole pH 7.5, 0.25% DM) and the protein was eluted with buffer ELU (200 mM NaCl, 20 mM HEPES pH 7, 10% Glycerol, 200 mM imidazole pH 7.5, 0.25% DM). Tag cleavage proceeded by addition of HRV 3C protease at a 3:1 molar ratio and dialysis against buffer DIA (200 mM NaCl, 20 mM HEPES pH7, 10% Glycerol, 0.25% DM) for at least 2 h at 4 °C. After application to NiNTA resin to remove the tag, the sample was concentrated by centrifugation using a 50 kDa molecular weight cutoff concentrator (Amicon) and further purified by size exclusion chromatography using a Superdex S200 column (GE Healthcare) pre-equilibrated with buffer SEC (200 mM NaCl, 20 mM HEPES pH 7, 0.25% DM). The peak fractions were pooled, concentrated, and used directly for further experiments.

### Cell culture

HEK293S GnTI^-^ and HEK293T cells were obtained from ATCC. HEK293S GnTI^-^ cells were grown in HyCell HyClone Trans Fx-H media supplemented with 1% FBS, 4 mM l-glutamine, 100 U/ml penicillin and 100 μg/ml streptomycin, and 1.5 g/l Kolliphor-188 at 37°C and 5% CO2. Adherent HEK293T cells were grown in DMEM-Glutamax, supplemented with 10% FBS, 100 U/ml penicillin and 100 μg/ml streptomycin. All cell lines were regularly tested for prevention of mycoplasma contamination.

### Expression screening of NRAT homologues

The DNA coding for NRATs from different plants was synthesized by Genscript and cloned into a pcDNA3.1 vector (Invitrogen) using the FX cloning technique (Geertsma and Dutzler, 2011) yielding a final construct where each NRAT1 contains a HRV 3C protease cleavage site, eGFP, a Myc tag and a streptavidin binding protein (SBP) attached to either its N- or C-terminus. Adherent HEK 293T cells (ATCC CRL-1573) were used for expression screening of plant NRAT1 homologues. Cells were grown on plates in DMEM-Glutamax High Glucose supplemented with 10% FBS, 100 U/ml penicillin and 100 μg/ml streptomycin. Cells were grown and split to 60-70% confluency 1 day prior to transfection. Cells were transiently transfected using polyethyleneimine (PEI) 25K. Briefly, for each 10 ml-plate, 10 μg of DNA in 500 μl of non-supplemented DMEM was mixed with 40 μg of PEI 25K in 500 μl of non-supplemented DMEM. After 15 min incubation, the transfection mixture was added to the plate together with 3 mM valproic acid. Cells were grown at 37 °C and 5% CO2 for 48-60 h and harvested by centrifugation at 1,000 g for 15 min. Pellets were resuspended in buffer LYS (200 mM NaCl, 20 mM HEPES pH7, 10% glycerol, 2 mM MgCl_2_, 40 μg/ml DNAse I, protease inhibitors (Roche Complete, EDTA-free) and 1% DDM). The lysate was incubated at 4 °C for 1.5 h under gentle agitation and clarified by centrifugation at 200,000 g for 15min. The clarified whole-cell extract was subjected to fluorescent size exclusion chromatography (Agilent HPLC) using a Superdex S200 column (GE Healthcare) and detected with a fluorescence detector set to λ_ex_= 489 nm / λ_em_= 510 nm. SiNRAT from *Setaria italica* (Uniprot KB K3YRE7) with a C-terminal tag was selected as the biochemically best behaving homologue based on the monodispersity of the eluted peak (Hattori et al., 2012).

### Expression and Purification of SiNRAT

A plasmid carrying the sequence coding for the NRAT from the plant *Setaria italica* (SiNRAT) (Uniprot KB K3YRE7) cloned into a pcDNA3.1 vector (Invitrogen) containing a HRV 3C protease cleavage site, eGFP, a Myc tag and a streptavidin binding protein (SBP) attached to its C-terminus was used for all further experiments. All mutants were generated using the QuickChange site-directed mutagenesis method (Agilent) (Table 2). SiNRAT was expressed by transient transfection using HEK293S GnTI^-^ cells diluted to 0.6-0.8.10^6^ cells/ml one day prior to transfection. DNA for transfection was purified from *E. coli* MC1061 cell line using the Nucleobond Xtra Maxi Kit (Macherey-Nagel). 1.3 μg DNA was used per 10^6^ cells. For that purpose, the DNA was diluted to 16 μg/ml in a volume of non-supplemented DMEM corresponding to 1/14 of the transfected cell culture volume. In parallel, PEI-max 40 kDA was diluted to 0.04 mg/ml in the same volume of non-supplemented DMEM. The two solutions were mixed and after 15-20 minutes incubation, the final transfection mixture was added to the suspension cells. Finally, the cell culture was supplemented with 3 mM valproic acid. After 48-60 h incubation, cells were harvested by centrifugation at 1,500g for 20 min and washed with PBS. All following steps of the purification were carried out at 4 °C. The cell pellets were resuspended in buffer RES (200 mM NaCl, 50 mM KPi pH 7.5, 10% Glycerol, 2 mM MgCl_2_, 40 μg/ml DNAse I and protease inhibitor (Roche Complete, EDTA-free)). The protein was extracted by adding n-dodecyl-β-D-maltopyranoside (DDM, Anatrace) to a final concentration of 2%. After an incubation of 2 h under gentle agitation, the lysate was clarified by centrifugation at 200,000 g for 30 min. The supernatant was loaded on Streptactin Superflow affinity resin (IBA Lifesciences) and incubated for 2 h under gentle agitation. The resin was washed with 25 CV of buffer W (200 mM NaCl, 20 mM HEPES pH 7, 10% glycerol, 0.04% DDM) and eluted with 20 ml of buffer ELU (buffer W supplemented with 5 mM D-desthiobiotin). The eGFP-Myc-SBP tag was removed by adding HRV 3C protease at a 1:3 molar ratio with SiNRAT and incubated on ice for 1 h. The protein was concentrated by centrifugation to 500 μl using a 50 kDa molecular weight cut-off concentrator (Amicon) and further purified by size exclusion chromatography on a Superdex S200 column (GE Healthcare) pre-equilibrated with buffer SEC (200 mM NaCl, 20 mM HEPES pH 7, 0.04% DDM).

### Isothermal titration calorimetry

A MicroCal ITC200 system (GE Healthcare) was used for ITC experiments. All titrations were performed at 6 °C in buffer SEC (200 mM NaCl, HEPES pH 7, 0.04% DDM). The syringe was filled with 1 mM AlCl3 and the titration was initiated by the sequential injection of 2 μl aliquots into the cell filled with SiNRAT or mutants at a concentration of 30 μM. Data were analyzed with the Origin ITC analysis package and fitted WT protein data fitted using the integrated two sides model. Each experiment was repeated with independently purified protein samples with similar results. Experiments with buffer not containing any protein were performed as controls.

### Reconstitution of SiNRAT and EcoDMT into proteoliposomes

SiNRAT and its mutants were reconstituted into detergent destabilized liposomes (Geertsma et al., 2008). The lipid mixture consisting of POPE, POPG and EggPC (Avanti Polar Lipids) at a weight ratio of 3:1:1 was washed with diethylether and dried under a nitrogen stream and by exsiccation overnight. The dried lipids were resuspended into 100 mM KCl and 20 mM HEPES pH 7. After 3 freeze-thaw cycles, the lipids were extruded through a 400 nm polycarbonate filter (Avestin, LiposoFast-BAsic) and aliquoted into samples at a concentration of 45 mg/ml. The extruded lipids were destabilized by addition of Triton X-100 and the protein was reconstituted at a protein to lipid ratio of 1:50. After several incubation steps with biobeads SM-2 (Biorad), the proteoliposomes were harvested the next day, resuspended in 100 mM KCl and 20 mM HEPES pH 7 and stored at −80 °C.

### Fluorescence-based substrate transport assays

The proteoliposomes were incubated with buffer IN (100 mM KCl, 20 mM HEPES pH 7, containing either 250 μM Calcein (Invitrogen), 100 μM Fura-2 (ThermoFischer Scientific), or 400 μM MagGreen (ThermoFischer Scientific) depending on the assayed ion). After 3 cycles of freeze-thawing, the liposomes were extruded through a 400 nm filter and the proteoliposomes were centrifuged and washed 3 times with buffer IN not containing any fluorophores. The proteoliposomes were subsequently diluted to 0.025 mg/ml in buffer OUT (100 mM NaCl, 20 mM HEPES pH 7) and aliquoted in 100 μl batches into a 96-well plate (ThermoFischer Scientific) and the fluorescence was measured every 4 seconds using a Tecan Infinite M1000 fluorimeter. An inside negative membrane potential was generated by addition of valinomycin to a final concentration of 100 nM. After 20 cycles of incubation, the substrate was added at different concentrations. The transport reaction was terminated by addition of calcimycin or ionomycin at a final concentration of 100 nM. The initial rate of transport was calculated by linear regression within the initial 100 seconds of transport after addition of the substrate. The calculated values were fitted to a Michaelis-Menten equation. Fura-2 was used to assay the transport of Ca^2+^. For detection, the ratio between calcium bound Fura-2 (λ_ex_= 340 nm; λ_em_= 510 nm) and unbound Fura-2 (λ_ex_= 380 nm; λ_em_= 510 nm) was monitored during kinetic measurements. Calcein was used to assay the transport of Mn^2+^ (λ_ex_= 492 nm; λ_em_= 518 nm) and Magnesium Green was to assay the transport of Mg^2+^ (λ_ex_= 506 nm; λ_em_= 531 nm).

For assaying coupled proton transport, the proteoliposomes were mixed with buffer 1 (100 mM KCl, 5 mM HEPES pH 7, 50 μM ACMA). After clarification by sonication, the proteoliposomes were diluted to a concentration of 0.15 mg/ml in buffer 2 (100 mM NaCl, 5 mM HEPES pH 7) and aliquoted into batches of 100 μl into a flat black 96-well plate (ThermoFischer Scientific). The fluorescence was measured every 4 seconds using a fluorimeter Tecan Infinite M1000 (λ_ex_= 412 nm; λ_em_= 482 nm). The transport reaction was initiated after addition of the substrate and valinomycin at a final concentration of 100 nM. The reaction was terminated by addition of CCCP at 100 nM. For experiments involving Ga^3+^, GaI3 was used. For compensation, NaI was added to reach a final I-concentration of 45 μM, which did not exert any effect on the measurement.

### Generation of nanobodies specific to SiNRAT

SiNRAT specific nanobodies were generated using a method described previously (Pardon et al., 2014). Briefly, an alpaca was immunized four times with 200 μg of detergent solubilized SiNRAT (in intervals of four weeks). Two days after the final injection, the peripheral blood lymphocytes were isolated, and the RNA total fraction was extracted and converted into cDNA by reverse transcription. The nanobody library was amplified and cloned into the phage display pDX vector (Invitrogen). After two rounds of phage display, ELISA assays were performed on the periplasmic extract of 198 individual clones. A single nanobody was identified. For all steps of phage display and ELISA, SiNRAT was chemically biotinylated using a EZ-Link™ NHS-PEG4 Biotinylation Kit (TermoFischer Scientific) and bound (Fairhead and Howarth, 2015) to neutravidin coated plates. The biotinylation was confirmed by total mass spectrometry analysis.

### Expression and purification of nanobodies and preparation of the SiNRAT-Nb1 complex

Nb1 was cloned into the pBXNPHM3 vector (Invitrogen) using the FX cloning technique. The expression construct contains the nanobody with a fusion of a pelB sequence, a 10-His tag, a maltose-binding protein (MBP) and a HRV 3C protease site to its N-terminus. The vector was transformed into *E. coli* MC1061 cells and a preculture was grown in TB-Gly medium overnight using ampicillin (100 μg/ml) as a selection marker in all expression cultures. The preculture was used for inoculation of TB-Gly medium at 1:100 volume ratio. Cells were grown to an OD600 of 0.7-0.9 and the expression was induced by addition of arabinose to a final concentration of 0.02% (w/v) for 4 h at 37 °C. Cells were harvested by centrifugation for 20 min at 5,000 g, resuspended in buffer A (200 mM KCl, 50 mM KPi pH 7.5, 10% Glycerol, 15 mM imidazole pH 7.5, 2 mM MgCl_2_, 1μM leupeptin, 1 μM pepstatin, 0.3 μM aprotinin, 1mM benzamidine, 1mM PMSF) and lysed using a high pressure cell lyser (Maximator HPL6). The purification was carried out at 4 °C. The lysate was cleared by centrifugation at 200,000 g for 30 min. The supernatant was loaded onto NiNTA resin and incubated for at least 1 h under gentle agitation. The resin was washed with 20 CV of buffer B (200 mM KCl, 20 mM Tris-HCl pH 8, 10% Glycerol, 50 mM imidazole, pH 7.5). The protein was eluted with buffer C (200 mM KCl, 20 mM Tris-HCl pH 8, 10% Glycerol, 300 mM imidazole). The tag was cleaved by addition of HRV 3C protease at a 3:1 molar ratio and dialyzed against buffer D (200 mM KCl, 20 mM Tris-HCl pH 8, 10% Glycerol) for at least 2 h at 4 °C. After application of NiNTA resin to remove the tag, the sample was concentrated by centrifugation using a 10 kDa molecular weight cut off concentrator (Amicon) and further purified by size exclusion chromatography using a Superdex S200 column (GE Healthcare) pre-equilibrated with buffer E (200 mM KCl, 20 mM Tris.HCl pH 8). The peak fractions were pooled, concentrated and flash-frozen and stored at −20 °C for subsequent experiments.

### Amphipol reconstitution of SiNRAT

For structure determination, the SiNRAT-Nb1 complex was reconstituted into amphipols. To ensure the stability of the proteins during this process, purified SiNRAT and Nb1 were mixed at a 1:2.5 molar ratio for 30 min prior to amphipol reconstitution. The SiNRAT-Nb1 complex and amphipol PMAL-C8 were mixed at a mass ratio of 1:5 and gently stirred overnight at 4 °C. DDM was removed by addition of a 100-fold excess (w/w) of biobeads SM-2 (Biorad) compared to protein for 4 h at 4 °C under gentle agitation. The biobeads were removed by filtration and the protein-amphipol mixture was concentrated before purification by size exclusion chromatography using a Superdex S200 column (GE Healthcare) pre-equilibrated with buffer SEC2 (200 mM NaCl, 20 mM HEPES pH 7). The protein peak fractions were pooled and concentrated to 2.7 mg/ml for cryoEM grid preparation.

### Cryo-EM sample preparation and data collection

For sample preparation for cryo-EM, 2.5 μl of the complex were applied to glow-discharged holey carbon grids (Quantifoil R1.2/1.3 Au 200 mesh). Samples were blotted for 2-4 s at 4 °C and 100% humidity. The grids were frozen in liquid propane-ethane mix using a Vitrobot Mark IV (Thermo Fisher Scientific). All datasets were collected on a 300 kV Titan Krios (ThermoFischer Scientific) with a 100 μm objective aperture and using a post-column BioQuantum energy filter with a 20 eV slit and a K3 direct electron detector (Gatan) in super-resolution mode. All datasets were recorded automatically using EPU2.9 (Thermo Fischer) with a defocus ranging from −1 to −2.4 μm, a magnification of 130,000x corresponding to a pixel size of 0.651 Å per pixel (0.3255 Å in super resolution mode) and an exposure of 1.01s (36 frames). Two datasets were collected for SiNRAT-Nb1 with respective total doses of 69.739 and 69.63 e^-^/Å^2^.

### Cryo-EM data processing

Datasets were processed using Cryosparc v3.2.0 (Punjani et al., 2017) following the same processing pipeline (Figure 2—figure supplement 2). All movies were subjected to motion correction using patch motion correction with a Fourier crop factor of 2 (pixel size of 0.651 Å/pix). After patch CTF estimation, high quality micrographs were identified based on relative ice thickness, CTF resolution and total full frame motion and micrographs not meeting the specified criteria were rejected. For the SiNRAT-Nb1 complex, 2D classes generated from the final map of rXkr9 in complex with Sb1^rXkr9^ (Straub et al., 2021) was used for template-driven particle picking on the whole dataset.

The 2D classes generated from particles selected from the entire dataset were extracted using a box size of 360 pix and down-sampled to 180 pixels (pixel size of 1.32 Å/pix). These particles were used for generation of 4 *ab initio* classes. Promising *ab initio* models were selected based on visual inspection and subjected to heterogenous refinement using one of the selected models as ‘template’ and an obviously bad model as decoy model. After several rounds of heterogenous refinement, the selected particles and models were subjected to non-uniform refinement (input model filtering to 8 Å) followed by local CTF refinement and another round of non-uniform refinement. Finally, the maps were sharpened using the sharpening tool from the Cryosparc package. The quality of the map was evaluated validated using 3DFSC (Tan et al., 2017) for FSC validation and local resolution estimation.

### Cryo-EM model building and refinement

Model building was performed in Coot (Emsley and Cowtan, 2004). Initially, the structure of ScaDMT (PDB 5M94) was rigidly fitted into the densities and served as template for map interpretation. The quality of the map allowed for the unambiguous assignment of residues 54-497. The structure of Nb16 (PDB 5M94) was used to build Nb1. The model was iteratively improved by real space refinement in PHENIX (Afonine et al., 2018) maintaining secondary structure constrains throughout. Figures were generated using ChimeraX (Pettersen et al., 2021) and Dino (http://www.dino3d.org). Surfaces were generated with MSMS (Sanner et al., 1996).

## Author Information

The authors declare no competing financial interests. Correspondence and requests for materials should be addressed to R.D. and C.M.

## Data availability

The cryo-EM density map of the SiNRAT-Nb1 complex and the coordinates for the atomic model will be made available upon publication.

## Acknowledgements

This research was supported by the Swiss National Science Foundation (SNF) through the National Centre of Competence in Research TransCure. We thank Dr. Marta Sawicka for input in cryo-EM and help during initial sample characterization. Nanobodies were generated with the help of the Nanobody Service Facility of UZH with the help of Dr. Sasa Stefanic. The assistance of Yvonne Neldner during library generation is acknowledged. The cryo-electron microscope and K3-camera were acquired with support of the Baugarten and Schwyzer-Winiker foundations and a Requip grant of the Swiss National Science Foundation. We thank Simona Sorrentino and the Center for Microscopy and Image Analysis (ZMB) of the University of Zurich for their support and access to the electron microscope. All members of the Dutzler lab are acknowledged for help in various stages of the project.

---

**Figure 1—figure supplement 1.**
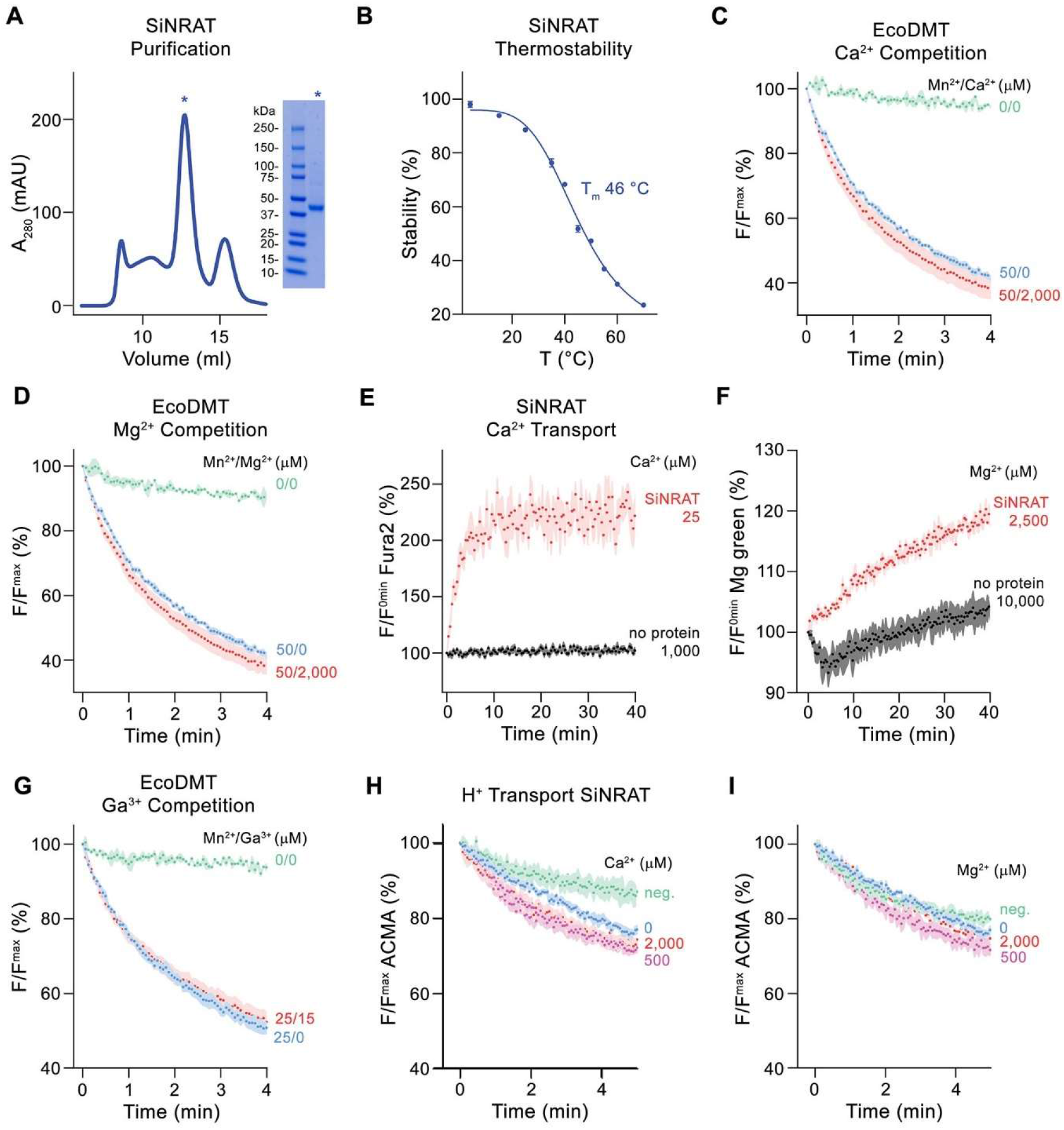
Purification and assay data. (**A**) Size exclusion chromatogram of purified SiNRAT. SDS PAGE gel of the purified peak fraction (asterisk) is shown right. (**B**) Thermal stability of SiNRAT assayed by fluorescence-based size-exclusion chromatography. Data show averages of 3 technical replicates, errors are s.e.m.. (**C**) Mn^2+^ transport in presence of Ca^2+^ (3 experiments for the condition with 50 μM Mn^2+^, 2 experiments for the condition with no substrate and 6 experiments for the condition with 50 μM Mn^2+^ and 2000 μM Ca^2+^, all from 2 independent reconstitutions) and (**D**) of Mg^2+^ (3 experiments for the condition with 50 μM Mn^2+^, 2 experiments for the condition with no substrate and 6 experiments for the condition with 50 μM Mn^2+^ and 2000 μM Mg^2+^, all from 2 independent reconstitutions) into proteoliposomes containing EcoDMT. (**E**) Comparison of Ca^2+^ import into proteoliposomes containing SiNRAT and protein-free liposomes assayed with the fluorophore Fura2 trapped inside the liposome (3 experiments from 3 independent reconstitutions for SiNRAT in presence of 25 μM Ca^2+^ and 8 experiments from 3 independent reconstitutions for empty liposomes in presence of 1000 μM Ca^2+^). (**F**) Comparison of Mg^2+^ import into proteoliposomes containing SiNRAT and protein-free liposomes assayed with the fluorophore Magnesium green (3 experiments from 3 independent reconstitutions for all conditions). (**G**) Mn^2+^ transport into proteoliposomes containing EcoDMT in presence of Ga^3+^ (3 experiments for the condition with 50 μM Mn^2+^, 2 experiments for the condition with no substrate and 6 experiments for the condition with 50 μM Mn^2+^ and 2000 μM Ca^2+^, all from 2 independent reconstitutions). **H**, **I** Experiments probing metal ion coupled H^+^ transport assayed with the fluorophore ACMA into proteoliposomes containing SiNRAT upon addition of Ca^2+^ (4 experiments from 3 independent reconstitutions for all conditions except 0.5 mM Ca^2+^ for which only 3 experiments were recorded) (**H**) and Mg^2+^ (4 experiments from 3 independent reconstitutions for all conditions except 0.5 mM Mg^2+^ for which only 3 experiments were recorded) (**I**). **A-G** and **J-L**, Panels show mean of indicated number of experiments, errors are s.e.m.. **C-I**, Fluorescence is normalized to the value after addition of substrate (t=0). Applied ion concentrations are indicated. Negative controls (neg.) refer to empty liposomes in presence of 2mM Ca^2+^ or 2mM Mg^2+^, respectively.

**Figure 2—figure supplement 1.**
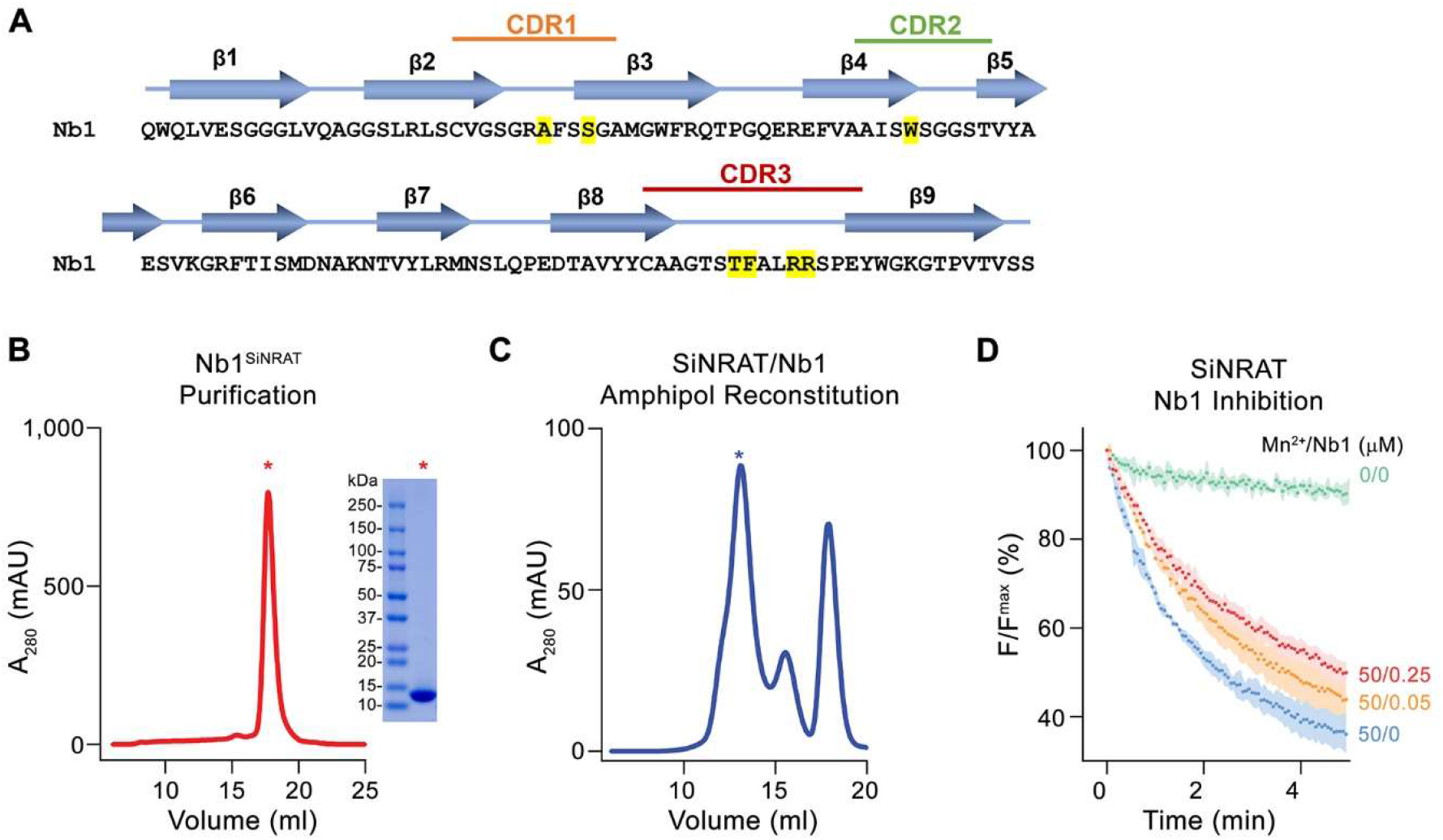
Nanobody characterization. (**A**). Sequence of Nb1 with secondary structure elements shown above. Complementary determining regions (CDR) are indicated and residues in contact with SiNRAT are highlighted in yellow. (**B**) Size exclusion chromatogram of purified Nb1. SDS PAGE gel of the purified peak fraction (asterisk) is shown on the right. (**C**) Size exclusion chromatogram of the purified SiNRAT-Nb1 complex reconstituted in amphipols. The peak fraction used for the preparation of cryo-EM grids is indicated by an asterisk. (**D**) Concentration-dependent inhibition of Mn^2+^ uptake into proteoliposomes containing SiNRAT upon addition of Nb1 to the outside. Data show mean of 4 experiments from 2 independent reconstitutions for all conditions except 0.25 μM Nb1 with 5 experiments, errors are s.e.m..

**Figure 2—figure supplement 2.**
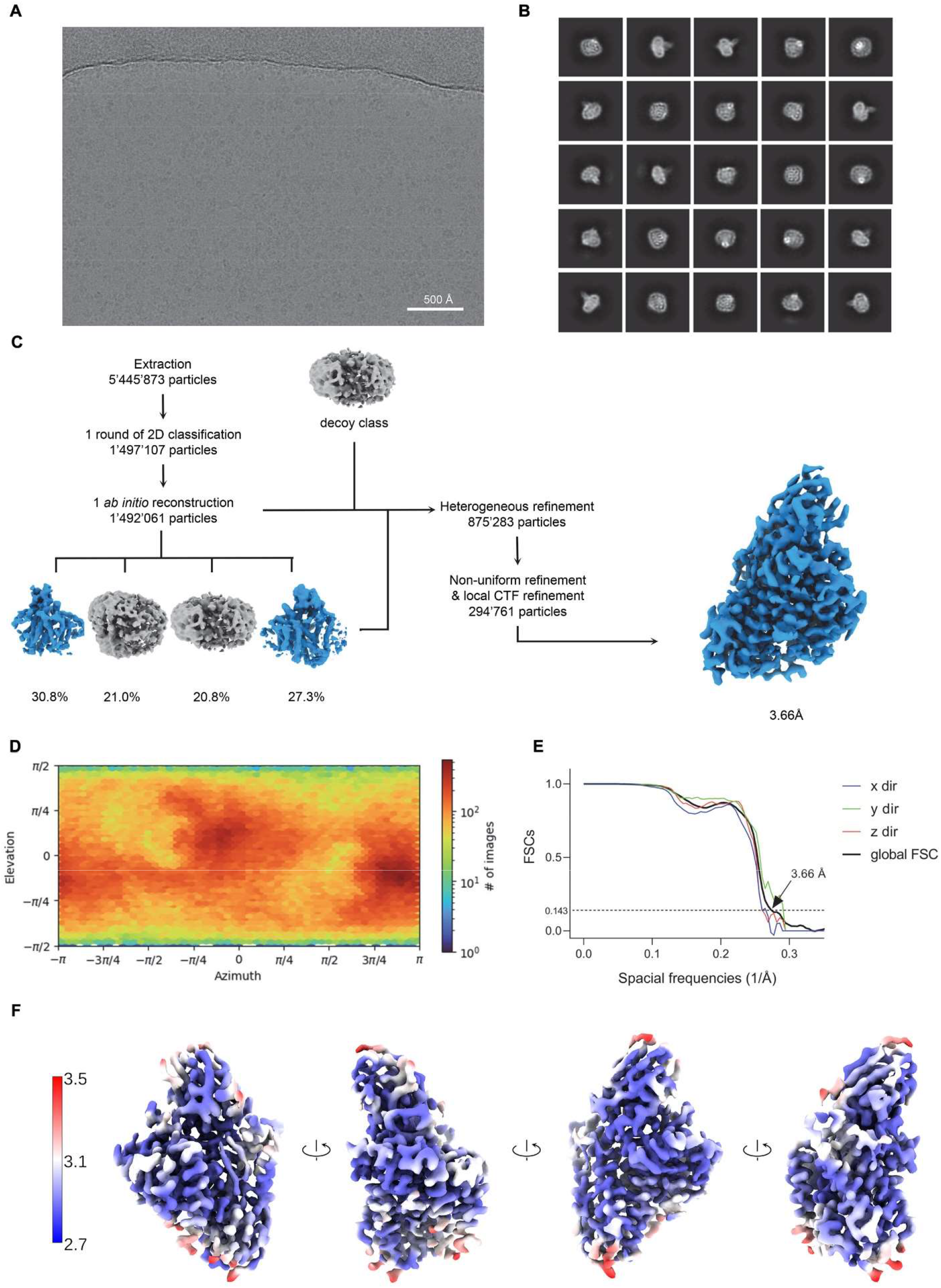
Cryo-EM reconstruction of the SiNRAT-Nb1 complex. (**A**) Representative micrograph (out of a total of 16,530 images) of the complex acquired with a Titan Krios G3i microscope equipped with a K3 camera. (**B**) 2D class averages of the SiNRAT-Nb1 dataset. (**C**) Data processing workflow. After extraction and one round of 2D classification, a single *ab initio* reconstruction with four classes was performed. The particle distribution is indicated in %. The two most promising classes, together with a decoy class were subjected to 9 rounds of heterogeneous refinement. Non-uniform (NU) refinement, local CTF-refinement and a second round of NU-refinement with C1 symmetry yielded a map at a resolution of 3.66 Å. (**D**) Angular distribution of particle orientations. The heatmap displays the number of particles for a given viewing angle. (**E**) Directional and global FSC plots. The global FSC is shown in black. Dashed line indicates 0.143 cut-off and the resolution at which the FSC curve drops below 0.143 is indicated. The directional FSC curves providing an estimation of anisotropy of the dataset are shown for directions x, y, and z. (**F**) Final 3D reconstruction of the SiNRAT-Nb1 complex in indicated orientations, colored according to the local resolution, estimated in cryoSPARC v.3.2.0.

**Figure 2—figure supplement 3.**
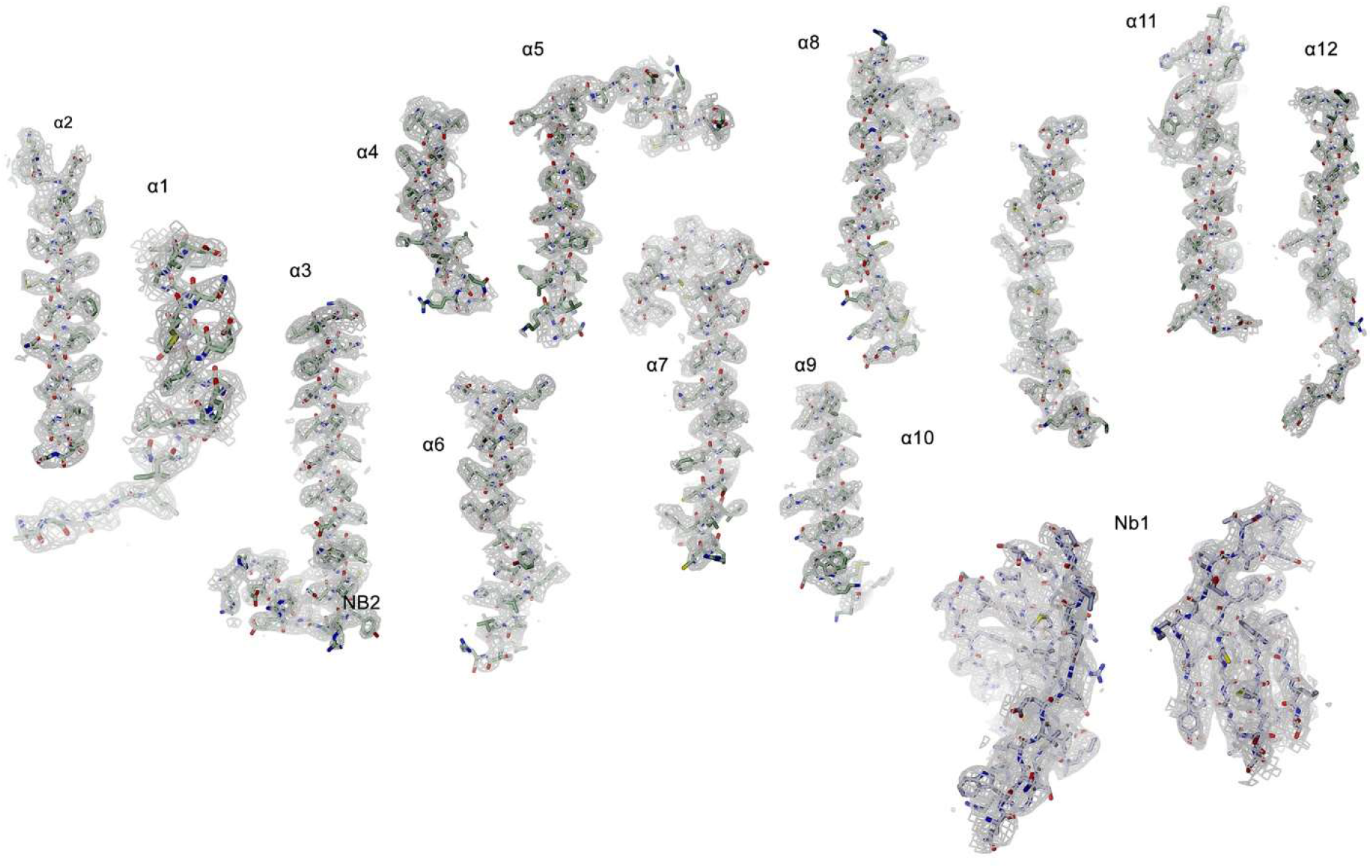
Cryo-EM density of the SiNRAT-Nb1 complex. Cryo-EM density at 3.66 Å (contoured at 6σ, gray mesh) is shown superimposed on indicated protein regions.

**Figure 2—figure supplement 4.**
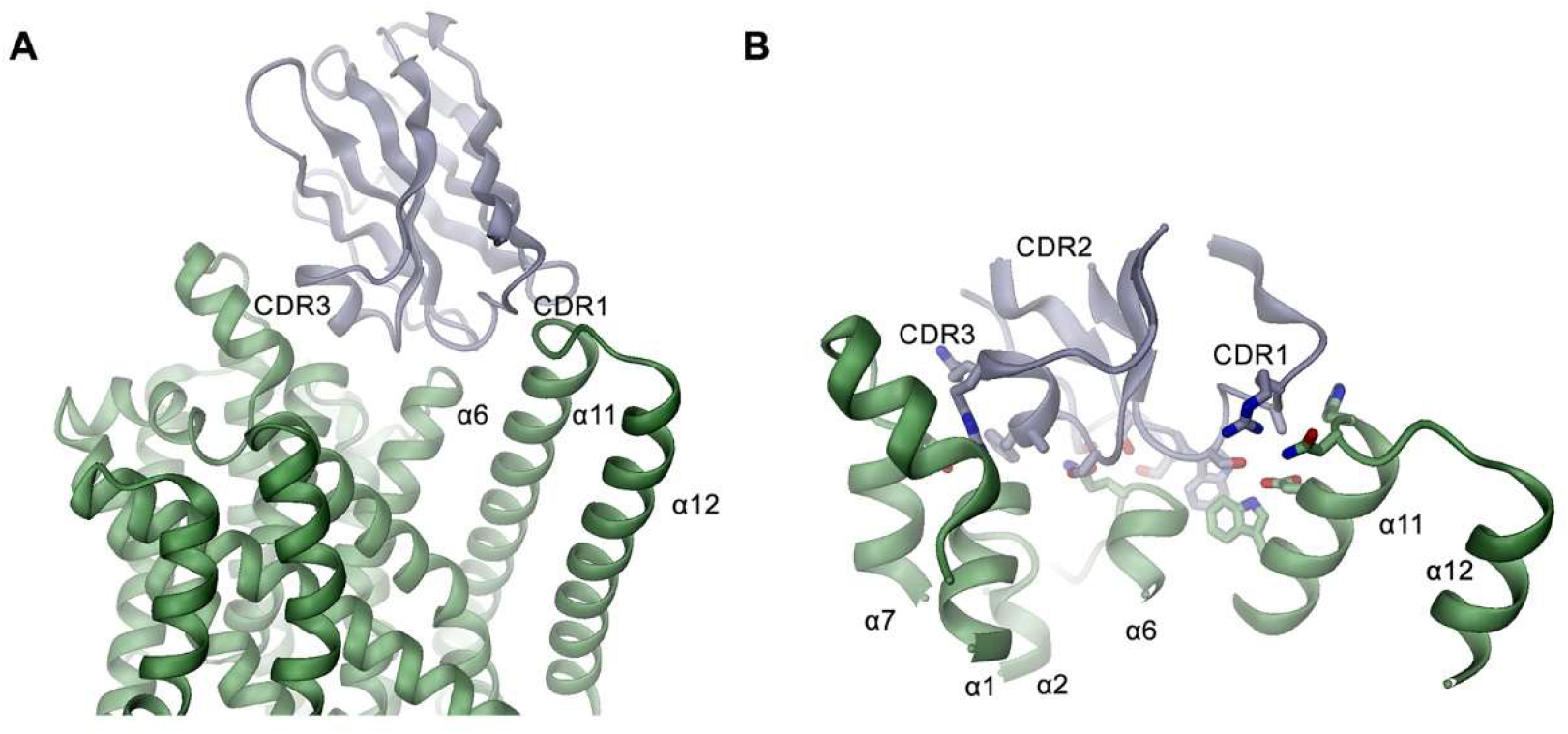
SiNRAT-Nb1 interactions. (A) Ribbon representation of the interaction region between SiNRAT and Nb1. (B) Closeup of the interaction region with interacting side chains shown as sticks.

